# BAR scaffolds drive membrane fission by crowding disordered domains

**DOI:** 10.1101/276147

**Authors:** Wilton T. Snead, Wade F. Zeno, Grace Kago, Ryan W. Perkins, J Blair Richter, Chi Zhao, Eileen M. Lafer, Jeanne C. Stachowiak

## Abstract

Cylindrical protein scaffolds are thought to stabilize membrane tubules, preventing membrane fission. In contrast, Snead et al. find that when scaffold proteins assemble, bulky disordered domains within them become acutely concentrated, generating steric pressure that destabilizes tubules, driving fission.

**Abstract:** Cellular membranes are continuously remodeled. The crescent-shaped bin-amphiphysinrvs (BAR) domains remodel membranes in multiple cellular pathways. Based on studies of BAR domains in isolation, the current paradigm is that they polymerize into cylindrical scaffolds that stabilize lipid tubules, preventing membrane fission. But in nature BAR domains are often part of multi-domain proteins that contain large intrinsically-disordered regions. Using in vitro and live cell assays, here we show that full-length BAR domain-containing proteins, rather than stabilizing membrane tubules, are instead surprisingly potent drivers of membrane fission. Specifically, when BAR scaffolds assemble at membrane surfaces, their bulky disordered domains become crowded, generating steric pressure that destabilizes lipid tubules. More broadly, we observe this behavior with BAR domains that have a range of curvatures. These data challenge the idea that cellular membranes adopt the curvature of BAR scaffolds, suggesting instead that the ability to concentrate disordered domains is the key requirement for membrane remodeling and fission by BAR domain-containing proteins.

## Introduction

Cellular membranes must undergo dynamic remodeling to facilitate essential cellular processes including formation of trafficking vesicles (Conner and Schmid, 2003), viral egress (Hurley et al., 2010), and cytokinesis (Mierzwa and Gerlich, 2014). Since membranes resist deformation (Helfrich, 1973), cells employ specialized protein machines to drive membrane remodeling (Zimmerberg and Kozlov, 2006). For example, the crescent-shaped, dimeric bin-amphiphysin-rvs (BAR) domains (Frost et al., 2009; Mim and Unger, 2012; Simunovic et al., 2015) polymerize into cylindrical scaffolds on membrane surfaces, forcing the underlying membrane to adopt the tubular geometry of the scaffold (Adam et al., 2015; Frost et al., 2008; Mim et al., 2012). This rigid scaffold is thought to stabilize membrane tubules, preventing their division into separate membrane compartments through the process of membrane fission (Boucrot et al., 2012).

Importantly, most studies on the membrane shaping behavior of BAR domains have examined the BAR domain in isolation, with significant portions of the protein removed. Examples include the N-terminal amphipathic helix BAR (N-BAR) domain of amphiphysin (Peter et al., 2004), the FCH BAR (F-BAR) domain of FCHo1/2 (Henne et al., 2010; Henne et al., 2007), the F-BAR domain of the neuronal migration protein srGAP2 (Guerrier et al., 2009), the F-BAR domains of the cytokinesis proteins Imp2 (McDonald et al., 2016) and Cdc15 (McDonald et al., 2015), and the inverted BAR (I-BAR) domains of MIM and ABBA (Mattila et al., 2007; Saarikangas et al., 2009), among others. These results have provided critical insight into the detailed geometry of BAR domain arrangement at membrane surfaces, helping to elucidate their mechanisms of membrane curvature sensing and induction. However, BAR domains do not typically exist in isolation in the cell, but rather as part of large, multi-domain proteins which also frequently contain long, intrinsically-disordered protein (IDP) domains of several hundred amino acids (Henne et al., 2010; Lee et al., 2007; Miele et al., 2004; Roberts-Galbraith and Gould, 2010; Wuertenberger and Groemping, 2015). How might these disordered domains influence the membrane remodeling behavior of BAR domains?

Recent work from our lab (Stachowiak et al., 2010; Stachowiak et al., 2012) and others (Bhagatji et al., 2009; Copic et al., 2012; Jiang et al., 2013; Vennema et al., 1996; Wu et al., 2014) has revealed that molecular crowding among proteins attached to membrane surfaces at high density generates steric pressure, which provides a potent force for membrane shaping. Further, previous work found that disordered domains, which occupy large footprints on the membrane surface in comparison to well-folded proteins of equal molecular weight (Hofmann et al., 2012), enhanced the efficiency of membrane bending and fission (Busch et al., 2015; Snead et al., 2017). However, a fundamental, unanswered question has limited the potential of protein crowding to explain membrane remodeling in cells - what brings bulky domains together to generate steric pressure? In particular, what keeps crowded proteins from simply diffusing away from one another, dissipating steric pressure and inhibiting membrane shaping? Proteins such as amphiphysin (Miele et al., 2004; Peter et al., 2004) and FCHo1/2 (Henne et al., 2010; Henne et al., 2007), which contain both scaffold-forming BAR domains and bulky disordered domains, present a possible solution to this problem. Specifically, the ability of BAR domains to form scaffolds has the potential to locally concentrate disordered domains such that steric pressure is amplified rather than dissipated.

Therefore, we set out to investigate the impact of disordered domains on the membrane remodeling ability of BAR proteins. To our surprise, we found that while isolated BAR domains formed stable membrane tubules, full-length amphiphysin and FCHo1 destabilized these tubules, leading to highly efficient membrane fission. These results challenge the paradigm that BAR scaffolds stabilize membrane tubes, and suggest instead that they act as templates that locally amplify steric pressure among disordered domains, leading to membrane fission.

## Results

### While the amphiphysin N-BAR domain stabilizes membrane tubules, full-length amphiphysin drives membrane fission

Amphiphysin is composed of an N-terminal, amphipathic helix BAR (N-BAR) domain, followed by an intrinsically disordered protein (IDP) domain of approximately 383 amino acids in humans, and a C-terminal SH3 domain (Miele et al., 2004; Owen et al., 2004; Owen et al., 1998; Peter et al., 2004) (Fig. 1A). To compare membrane bending by full-length amphiphysin (Amph-FL) to the N-BAR domain alone, we first examined the effects of each protein on giant unilamellar vesicles (GUVs). These experiments revealed that both the N-BAR domain and Amph-FL drove potent membrane bending, forming mobile, diffraction-limited tubules that extended from the GUV surface (Fig. S1A, Movies S1 and S2). These GUVs often collapsed or broke apart into smaller tubules and fragments (Movies S3 and S4), suggesting that lipid tubule formation may not have been the endpoint of the membrane remodeling process.

**Figure 1.**
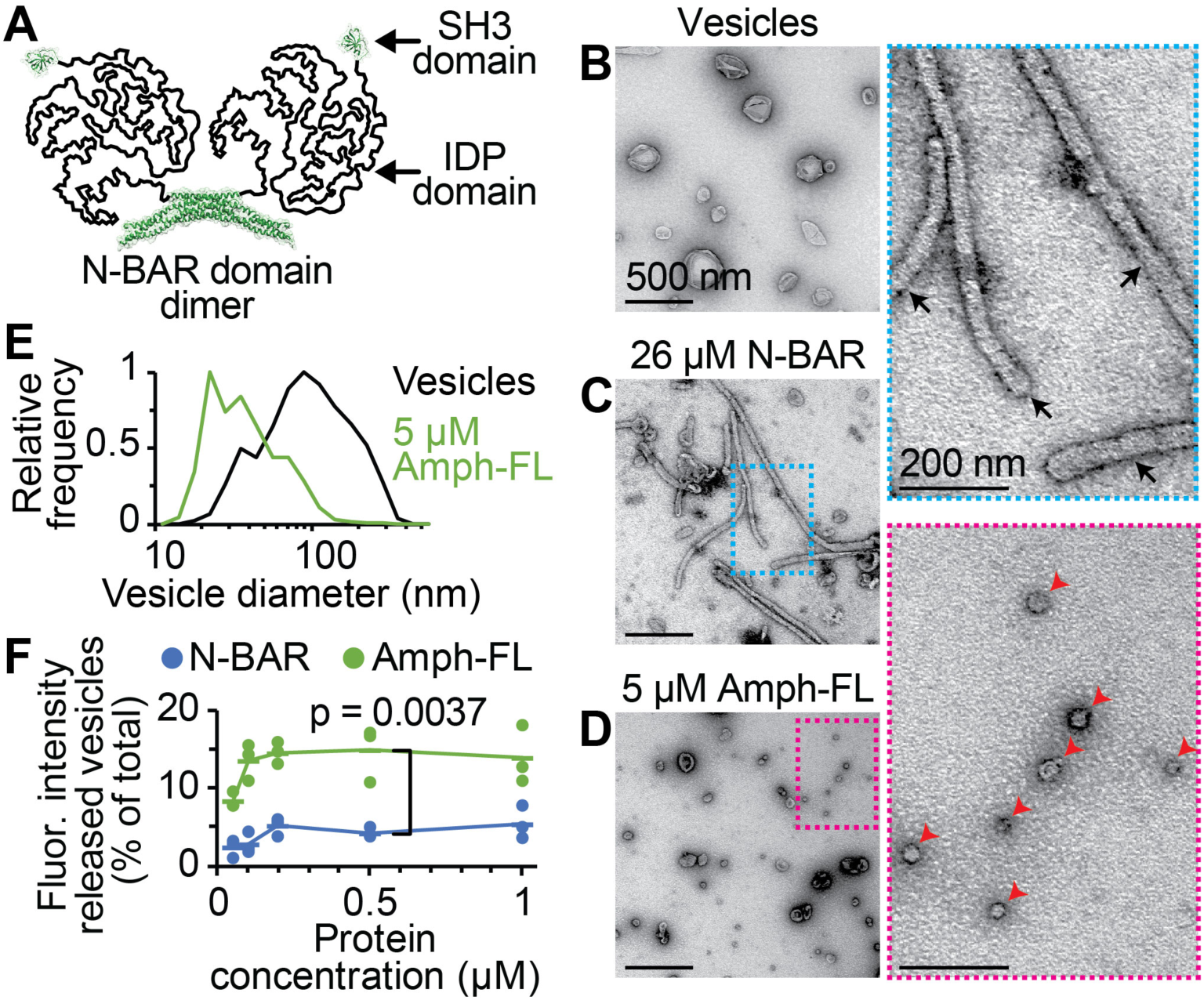
Amphiphysin drives membrane fission, while the N-BAR domain stabilizes membrane tubules. Membrane composition for vesicles in TEM: 80 mol% DOPC, 5 mol% PtdIns(4,5)P_2_, 15 mol% DOPS. SUPER template membrane composition: 79 mol% DOPC, 5 mol% PtdIns(4,5)P_2_, 15 mol% DOPS, 1 mol% Texas Red-DHPE. (**A**) Schematic of full-length amphiphysin (Amph-FL) dimer. BAR domain: PDB 4ATM. SH3 domain: PDB 1BB9. (**B-D**) Negative stain TEM micrographs of 200 nm extruded vesicles (**B**) before exposure to protein, (**C**) after exposure to 26 μM N-BAR, and (**D**) after exposure to 5 μM Amph-FL. Dashed boxes indicate zoomed regions to the right. Black arrows indicate membrane tubules, red arrowheads indicate fission vesicles. (**E**) Histograms of vesicle diameters measured from electron micrographs. Vesicles alone: n = 1,302 vesicles. 5 μM Amph-FL: n = 1,071 vesicles. (**F**) Membrane release from SUPER templates, measured as Texas Red signal present in the supernatant after sedimentation of the SUPER templates. Membrane release in the absence of protein was measured and subtracted as background. Dots indicate data and lines indicate mean, n = 3 independent experiments. P-value: one-tailed, unpaired Student’s t-test. Scale bars in (B-D): 500 nm. Zoomed region scale bars: 200 nm. See also Fig. S1 and Movies S1-S4.

To directly visualize the morphology of membranes at the end of remodeling, we utilized negative stain transmission electron microscopy (TEM) to resolve membrane structures below the optical diffraction limit. As expected from previous findings (Gallop et al., 2006; Peter et al., 2004), the N-BAR domain transformed vesicles that had an average initial diameter of 200 nm into long tubules with average outer diameter 44±6 nm s.d. (Fig. 1B,C and S1B,C). In contrast, Amph-FL did not drive appreciable membrane tubule formation in TEM experiments. Rather, Amph-FL divided the vesicles of initially 200 nm diameter into a population of highly curved vesicles with a peak diameter centered near 22 nm (Fig. 1D,E and S1D). This result suggests that Amph-FL is capable of driving membrane fission, a more energetically demanding process than membrane tubule formation (Campelo et al., 2012).

### Full-length amphiphysin generates highly curved fission products

To better understand the ability of amphiphysin to drive membrane fission, we compared N-BAR and Amph-FL in two additional assays of membrane fission. In the first of these experiments, we used supported bilayers with extra membrane reservoir (SUPER templates), which are glass beads surrounded by a low-tension membrane. Exposure of SUPER templates to fission-driving proteins results in measurable membrane release from the beads (Liu et al., 2011; Neumann et al., 2013; Pucadyil and Schmid, 2008). SUPER template experiments revealed that while both N-BAR and Amph-FL drove membrane release in the concentration range of 50-1,000 nM, Amph-FL drove more than twice as much (2.6 to 4.7-fold greater) membrane release at each concentration (Fig. 1F). Notably, membrane release does not directly imply efficient membrane fission, as both vesicles and lipid tubules can be shed from SUPER templates. Therefore, we next employed a tethered vesicle assay to quantify the distributions of fission vesicle diameters over a range of protein concentrations (Snead et al., 2017). Specifically, we tethered fluorescent vesicles to a coverslip passivated with PEG and PEG-biotin (Fig. 2A). Vesicles in these experiments contained a biotinylated lipid, which facilitated tethering to the substrate via binding to neutravidin. Vesicles also contained the fluorescent lipid Oregon Green 488-DHPE, which we used to quantify the brightness of each vesicle after imaging in confocal fluorescence microscopy (Aguet et al., 2013) (Fig. 2B). We then converted the resulting distributions of vesicle brightness to approximate distributions of vesicle diameter by calibrating against the initial vesicle diameter distribution measured using dynamic light scattering (see methods).

**Figure 2.**
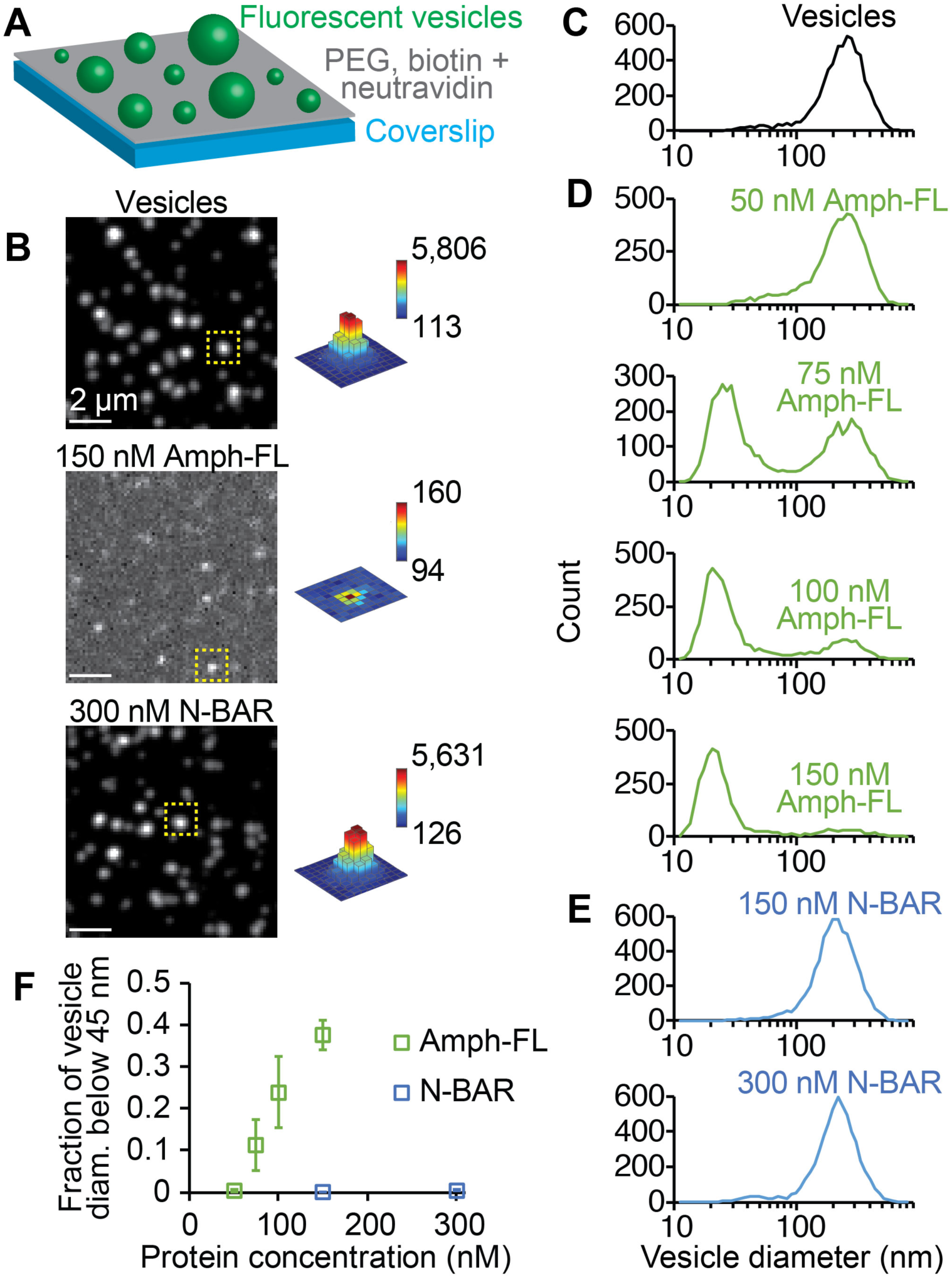
Full-length amphiphysin produces highly curved fission products. Tethered vesicle composition: 76 mol% DOPC, 5 mol% PtdIns(4,5)P_2_, 15 mol% DOPS, 2 mol% Oregon Green 488-DHPE, 2 mol% DP-EG10-biotin. (**A**) Schematic of tethered vesicle fission experiment. (**B**) Representative spinning disc confocal micrographs of tethered vesicles before exposure to protein (top), after exposure to 150 nM Amph-FL (middle), and after exposure to 300 nM N-BAR (bottom). Contrast settings in top and bottom images are the same while contrast in middle image is adjusted to clearly show vesicle puncta. Dashed yellow boxes indicate puncta intensity profiles on right, where bar heights are all scaled between 90 and 6,000 brightness units while each color map corresponds to specified intensity range. (**C-E**) Distributions of vesicle diameter measured by tethered vesicle assay (**C**) before exposure to protein, (**D**) after exposure to Amph-FL at 50, 75, 100, and 150 nM, and (**E**) after exposure to N-BAR at 150 and 300 nM. (**F**) Summary of tethered vesicle fission data, expressed as the ratio of the distribution area below 45 nm diameter to the total distribution area. Markers represent mean ± first s.d., n = 3 independent experiments. Scale bars in (B): 2 μm. See also Fig. S1-S3.

Using this assay, we found that Amph-FL in the concentration range of 50-150 nM transformed vesicles with an average initial diameter of 200 nm (Fig. 2B,C) to a population of high curvature fission products (Fig. 2B,D) with a median diameter of 22 nm at 150 nM, in agreement with results from TEM (Fig. 1D,E). The proportion of vesicles that fell within the high curvature group (diameters below approximately 45 nm) increased with increasing protein concentration, from less than 1% at 50 nM to approximately 38% at 150 nM (Fig. 2F). In contrast, the N-BAR domain did not drive fission, even at higher protein concentrations (Fig. 2B, E, and F). While lipid tubules were not apparent in tethered vesicle experiments with N-BAR (Fig. 2B), tubules were observed at micromolar N-BAR concentrations (Fig. S1E), consistent with the tubules formed in TEM experiments (Fig. 1C). Notably, higher concentrations of Amph-FL were required to observe fission in TEM experiments compared to tethered vesicle experiments. This increase is due to the high lipid concentration used in TEM experiments (approximately 100-fold greater than tethered vesicle experiments), which is necessary to achieve an adequate density of lipid structures for TEM (see methods). Taken together, our results from electron microscopy and tethered vesicle experiments confirm that Amph-FL is a potent driver of membrane fission, while the isolated N-BAR domain is only capable of forming membrane tubules.

### The ability of full-length amphiphysin to drive membrane fission does not arise from greater membrane binding affinity in comparison to isolated N-BAR

How can we explain the ability of full-length amphiphysin to drive membrane fission? One possible explanation could be that the full-length protein may bind more strongly to membrane surfaces compared to the N-BAR domain alone, resulting in more potent membrane remodeling. To examine this possibility, we utilized a tethered vesicle assay similar to the experiments described above to quantify the relative extent of protein-membrane binding. In this assay proteins were labeled with Atto 594 dye, while vesicles contained the lipid Oregon Green 488-DHPE. We quantified vesicle diameter as described above, and used measurements of single fluorophore brightness to quantify the number of bound proteins per vesicle (Snead et al., 2017) (see methods). From these measurements we determined the density of membrane-bound proteins, which increased with increasing protein concentration in solution (Fig. S2A-C). These experiments revealed that Amph-FL and N-BAR reached similar number densities of membrane-bound protein within the concentration range of 5-25 nM, indicating that Amph-FL and the isolated N-BAR domain bind with similar affinity to membranes (Fig. S2D,E). Specifically, these results suggest that the disordered domain did not significantly enhance protein-lipid and protein-protein interactions, either of which would be expected to increase the density of membrane-bound protein. Therefore, the ability of Amph-FL to drive membrane fission cannot be explained by differences in membrane recruitment.

### The disordered domain of amphiphysin drives membrane fission on its own, but requires higher protein concentration compared to full-length amphiphysin

Another possible explanation for the ability of amphiphysin to drive membrane fission is that its substantial disordered domain (residues 240-622) generates steric pressure that promotes fission, in line with previous studies on other membrane-bound disordered domains (Busch et al., 2015; Snead et al., 2017). If so, the isolated disordered domain should be able to drive membrane fission when bound to membrane surfaces at sufficient density. To test this idea, we purified the disordered domain of amphiphysin, lacking the C-terminal SH3 domain (Amph CTD ∆SH3, Fig. 3A). We first performed fluorescence correlation spectroscopy (FCS) measurements to quantify the hydrodynamic radius of this domain. Specifically, we calibrated the diffusion time of Amph CTD ∆SH3 against the diffusion times of two protein standards with known hydrodynamic radii, transferrin (Hall et al., 2002) and the C-terminal domain of AP180 (AP180 CTD) (Busch et al., 2015). These experiments yielded an approximate hydrodynamic radius for Amph CTD ∆SH3 of 4 nm (Fig. S2G-I, see methods), which corresponds to a radius of gyration of approximately 5 nm (Hofmann et al., 2012; Sherman and Haran, 2006), comparable to other disordered domains of similar molecular weight (Busch et al., 2015; Kalthoff et al., 2002). FCS experiments also showed that the size of Amph CTD ∆SH3 varied with the concentration of monovalent salt in the buffer (Fig. S2J), consistent with the known sensitivity of highly-charged disordered proteins to changes in ionic strength (Srinivasan et al., 2014). We next performed tethered vesicle experiments to assess membrane fission by Amph CTD ∆SH3. The protein included an N-terminal hexa-histidine (6his) tag to facilitate binding to membranes by the lipid DOGS-NTA-Ni (Fig. 3A). Experiments revealed that Amph CTD ∆SH3 indeed drove the formation of highly curved fission products (Fig. 3B,C). However, 100-fold greater concentration of Amph CTD ∆SH3 in solution (10 μM) was required to generate fission products of similar curvature to those observed with Amph-FL (100 nM) (see Fig. 2D,F).

**Figure 3.**
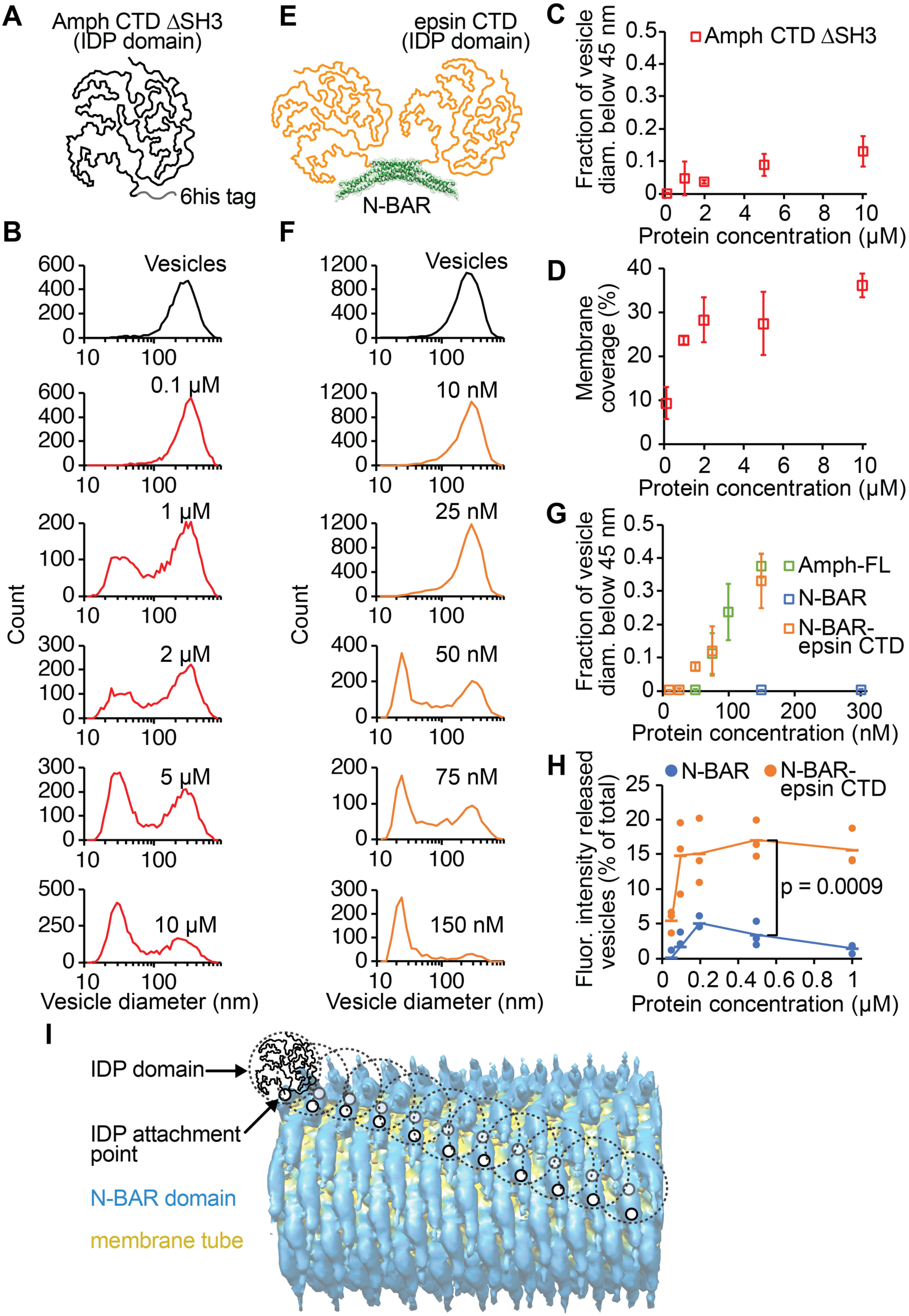
The disordered domain of amphiphysin alone drives membrane fission, but the N-BAR scaffold substantially enhances fission efficiency. Membrane composition in Amph CTD ∆SH3 tethered vesicle experiments: 76 mol% DOPC, 20 mol% DOGS-NTA-Ni, 2 mol% Oregon Green 488-DHPE, 2 mol% DP-EG10-biotin. In tethered vesicle experiments with N-BAR-epsin CTD, DOGS-NTA-Ni was replaced with 5 mol% PtdIns(4,5)P_2_ and 15 mol% DOPS. SUPER template membrane composition: 79 mol% DOPC, 5 mol% PtdIns(4,5)P_2_, 15 mol% DOPS, 1 mol% Texas Red-DHPE. (**A**) Schematic of Amph CTD ∆SH3. (**B**) Tethered vesicle fission experiments show that Amph CTD ∆SH3 forms highly curved fission products. (**C**) Summary of data from tethered vesicle fission experiments with Amph CTD ∆SH3 expressed as the ratio of the distribution area below 45 nm to the total distribution area (compare to Fig. 2F). (**D**) Coverage of the membrane surface by Amph CTD ∆SH3 for each concentration at which fission was measured. (**E**) Schematic of N-BAR-epsin CTD chimera dimer. (**F**) Tethered vesicle fission measurements show that N-BAR-epsin CTD generates highly curved fission vesicle populations over the concentration range of 10-150 nM, similar to Amph-FL (compare to Fig. 2D). (**G**) Summary of data from tethered vesicle fission experiments with N-BAR-epsin CTD, expressed as the ratio of the distribution area below 45 nm to the total distribution area. Amph-FL and N-BAR data from Fig. 2F. (**H**) SUPER template membrane shedding experiments show that N-BAR-epsin CTD drives greater membrane release compared to N-BAR (compare to Fig. 1F). Dots indicate data and lines indicate mean, n = 3 independent experiments. P-value: one-tailed, unpaired Student’s t-test. Markers in (C), (D), and (G) represent mean ± first s.d., n = 3 independent experiments. (**I**) Schematic of the N-BAR scaffold (EMDB 3192) (Adam et al., 2015) with attachment points of some of the disordered domains marked (two per N-BAR dimer). Dashed circles indicate approximate volumes occupied by undeformed disordered domains. See also Fig. S2 and S3.

Membrane binding experiments with fluorescently-labeled Amph CTD ∆SH3 showed that when 10 μM of protein was present in solution, the protein covered approximately 40% of the membrane surface (Fig. 3D and S2K-M). At this coverage, steric pressure from protein crowding is expected to be sufficient to overcome the energetic barrier to membrane fission (Snead et al., 2017). Therefore, the requirement for a high solution concentration of Amph CTD ∆SH3 reflects the conditions necessary to promote crowded binding to the membrane surface. In contrast, the ability of Amph-FL to drive fission at much lower concentrations likely arises from polymerization of the BAR domain scaffold, which enables multivalent membrane binding (Simunovic et al., 2016; Sorre et al., 2012). In line with this thinking, we found that 100 nM Amph-FL, a concentration which drove potent fission (Fig. 2D,F), covered greater than 70% of the membrane surface (Fig. S2F). This coverage is significantly higher than has been observed for non-assembling proteins (Feder, 1980; Snead et al., 2017). Collectively, these results suggest that the ability of amphiphysin to drive efficient membrane fission arises from a collaboration between its N-BAR domain and disordered domain.

### A chimera consisting of N-BAR fused to an alternative disordered domain drives fission with similar efficiency to wild-type amphiphysin

Experiments comparing membrane remodeling by Amph-FL and N-BAR imply that assembly of the N-BAR scaffold at the membrane surface facilitates local crowding of the bulky disordered domains in Amph-FL. This reasoning implies that any bulky disordered domain that is brought to the membrane surface by a BAR scaffold should be capable of driving membrane fission. To test this prediction, we created a chimera consisting of the amphiphysin N-BAR domain fused to the C-terminal disordered domain of rat epsin1 (N-BAR-epsin CTD, Fig. 3E). Importantly, the disordered domain of epsin1 is comparable to the disordered domain of amphiphysin in terms of length (432 versus 383 amino acids, respectively) as well as hydrodynamic radius (Busch et al., 2015). Tethered vesicle fission experiments revealed that N-BAR-epsin CTD generated highly curved fission products from vesicles with an initial average diameter of 200 nm within a similar range of protein concentrations to Amph-FL (Fig. 3F, compare to Fig. 2D). Further, both N-BAR-epsin CTD and Amph-FL produced a very similar fraction of fission products with diameters below 45 nm at equivalent concentrations in solution (Fig. 3G). Finally, SUPER template membrane shedding experiments revealed that in the concentration range of 50-1,000 nM, N-BAR-epsin CTD drove greater membrane release compared to the isolated N-BAR domain (Fig. 3H), similar to the results of SUPER template experiments comparing N-BAR and Amph-FL (Fig. 1F). These findings illustrate the ability of N-BAR scaffolds to promote membrane fission by crowding arbitrary disordered domains at membrane surfaces.

How does crowding among disordered domains overcome the ability of BAR scaffolds to stabilize lipid tubules? One explanation is that steric pressure among the bulky disordered domains of Amph-FL inhibits the assembly of a long-range N-BAR scaffold, which, if allowed to form, would inhibit fission. In support of this hypothesis, when Amph-FL reached over 70% surface coverage as described above (Fig. S2F), the underlying N-BAR domain covered only about 16% of the membrane, based on membrane footprints for Amph-FL and N-BAR of 79 and 16.5 nm^2^ per monomer, respectively (see methods). This coverage is significantly lower than expected for a fully-assembled N-BAR scaffold, which approaches complete coverage (Adam et al., 2015; Mim et al., 2012). Furthermore, the volume available per amphiphysin disordered domain above the N-BAR scaffold is only about 50% of the volume that each domain would be expected to occupy in solution, based on its radius of gyration (Fig. 3I, see calculation in methods). Therefore, the disordered domains would be required to compress substantially to fit around a fully-assembled N-BAR scaffold. Previous work has shown that substantial compression of disordered domains is energetically costly, likely exceeding the cost of membrane deformation (Busch et al., 2015). Collectively, these arguments suggest that the presence of amphiphysin’s bulky disordered domains inhibits assembly of long-range N-BAR scaffolds.

Interestingly, previous structural studies using cryo-electron microscopy have reported limited observations of tubular N-BAR scaffolds formed from full-length amphiphysin (Adam et al., 2015; Mim et al., 2012). These structures have been observed on membrane substrates containing a majority of negatively-charged lipids, which are thought to provide a strong electrostatic driving force for scaffold assembly (Adam et al., 2015; Mim et al., 2012). Therefore, we performed tethered vesicle fission experiments using a similar membrane composition (Fig. S3). Here, the onset of membrane fission occurred at somewhat higher Amph-FL concentration, 350 nM (Fig. S3) in comparison to 75 nM (Fig. 2B-F). These results demonstrate that high concentrations of negatively charged lipids cannot prevent membrane fission as protein concentration increases. Indeed, the cryo-electron microscopy studies cited above suggest that long-range scaffolds assembled from full-length protein were more rare (Mim et al., 2012). Moreover, these studies employed buffers that lacked small monovalent ions (Adam et al., 2015; Mim et al., 2012), conditions known to favor extension of disordered domains owing to reduced electrostatic screening (Srinivasan et al., 2014). This environment likely enabled the disordered domains to pack more efficiently around the scaffold, in line with the needle-like densities seen protruding from the N-BAR scaffold (Adam et al., 2015).

### Disordered domains inhibit tubule formation by BAR scaffolds in live cells

We next sought to evaluate the influence of disordered domains on the membrane remodeling behavior of BAR domains in live mammalian cells. Multiple previous studies have established that overexpression of BAR domains leads to formation of lipid tubules in the cytosol that are coated by BAR scaffolds (Boucrot et al., 2012; Frost et al., 2008; McDonald et al., 2015; Peter et al., 2004). We first replicated these findings by overexpressing the N-BAR domain of human amphiphysin tagged with mCherry (Fig. 4A) in retinal pigmented epithelial (RPE) cells. We found that N-BAR generated a dense network of tubules inside the cells (Fig. 4B), in agreement with previous findings (Peter et al., 2004). The number of tubules per cell increased with the expression level of N-BAR (Fig. 4C), and co-localized with a plasma membrane stain (Fig. S4A), indicating that many of the tubules originated from the plasma membrane as previously observed (McDonald et al., 2015; McDonald et al., 2016). In contrast, overexpression of Amph-FL tagged with mCherry (Fig. 4A) led to significantly fewer tubules per cell compared to N-BAR (Fig. 4 B-D), suggesting that the disordered domain of Amph-FL inhibited tubule formation. Notably, Amph-FL interacts with the clathrin adaptor network and may therefore recruit other membrane remodeling proteins. As such, it is unclear whether the lack of stable tubules in Amph-FL-expressing cells arose from the disordered domain or from other binding partners recruited by Amph-FL. To distinguish between these possibilities, we created a chimera of N-BAR fused to an alternative disordered domain, from the neuronal protein neurofilament-M (N-BAR-NfM CTD), tagged with mCherry (Fig. 4A). The disordered C-terminal domain of neurofilament-M acts as an entropic brush, radiating outward along the length of neurofilaments and sterically repelling neighboring disordered domains to control axon diameter (Brown and Hoh, 1997; Kumar et al., 2002). The neurofilament-M disordered domain is similar in length to that of amphiphysin (438 versus 383 amino acids, respectively) but is not involved in endocytosis and therefore contains no binding domains for endocytic proteins. Overexpressing N-BAR-NfM CTD in RPE cells resulted in a similar phenotype to Amph-FL, in which the number of tubules per cell was significantly reduced compared to the isolated N-BAR domain (Fig. 4 B-D). Furthermore, while tubules in N-BAR-expressing cells had an average length of 6.0±0.2 μm s.e.m., tubule lengths in cells expressing Amph-FL and N-BAR-NfM CTD were significantly shorter, 3.5±0.1 and 3.6±0.1 μm s.e.m., respectively (Fig. 4E).

**Figure 4.**
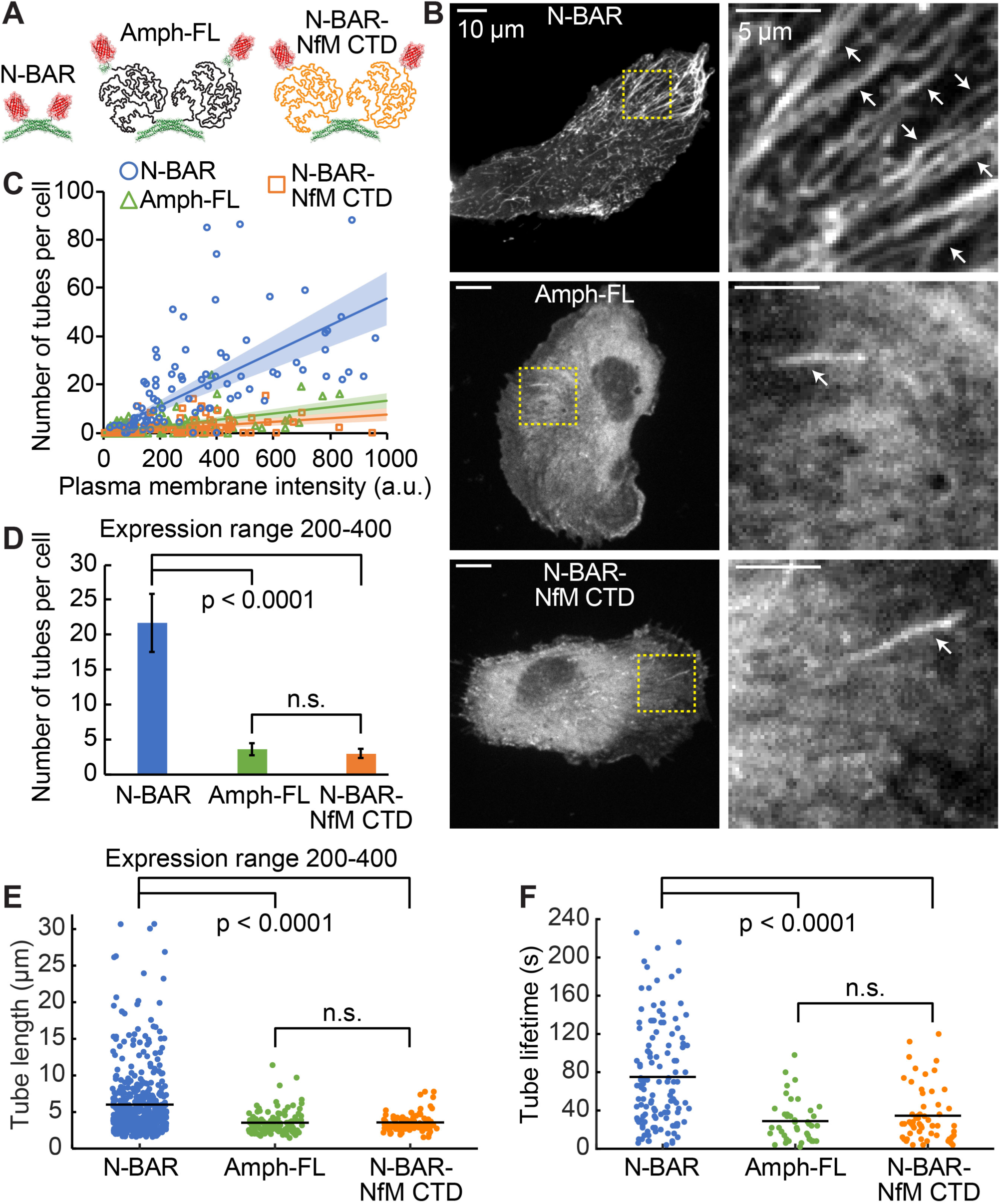
Disordered domains disrupt N-BAR mediated membrane tubulation in live cells. (**A**) Schematic of mCherry (PDB 2H5Q) fusion constructs expressed in cells. (**B**) Confocal images of RPE cells expressing N-BAR (top), Amph-FL (middle), and N-BAR-NfM CTD (bottom). Yellow dashed boxes indicate zoomed regions to the right. White arrows indicate tubules. All cells are within the same range of protein expression level used for quantification in (D) and (E). (**C**) Number of tubes per cell as a function of protein expression level, quantified as the background-subtracted protein intensity at the plasma membrane (see methods). Lines indicate linear regression with y-intercept set to 0. Shaded regions indicate 99% confidence interval. Line color matches the respective marker color. n > 90 cells per condition from two independent transfections. (**D**) Number of tubes per cell within the expression level range of 200-400 brightness units. Bars indicate mean ± s.e.m., n > 20 cells per condition. (**E**) Length of tubes in cells within the expression level range of 200-400 brightness units. Points indicate data, black lines indicate means. n > 80 tubes per condition. (**F**) Lifetime of tubes in cells measured from time-lapse TIRF movies (see methods). Points indicate data, black lines indicate means. n > 40 tubes per condition. All p-values: two-tailed, unpaired Student’s t-tests. Scale bars in (B): 10 μm. Zoomed region scale bars: 5 μm. See also Fig. S4 and Movie S5.

Time-lapse imaging of live cells revealed that the tubules formed by Amph-FL and N-BAR-NfM CTD were more transient in comparison to isolated N-BAR (Fig. 4F and Movie S5). Specifically, the tubules in cells expressing N-BAR had an average lifetime of approximately 75±5 s s.e.m., whereas tubule lifetime was significantly shorter in cells expressing Amph-FL and N-BAR-NfM CTD, approximately 29±3 and 35±4 s s.e.m., respectively (Fig. 4F). The tubules formed by N-BAR also had greater fluorescence intensity in the protein channel relative to the local background in comparison to Amph-FL and N-BAR-NfM CTD (Fig. S4B,C). This finding indicates that the disordered domains of Amph-FL and N-BAR-NfM CTD did not promote tubule fission by enhancing protein binding to the membrane surface. Collectively, results from experiments in live cells indicate that bulky disordered domains are capable of disrupting the formation of stable tubules scaffolded by BAR domains, similar to observations in vitro (Fig. 1C,D). The disordered domains may have inhibited tubule formation by driving membrane fission, though future work is needed to test the role of disordered domains in physiological fission events.

### Crowding among disordered domains opposes the ability of I-BAR scaffolds to drive inward membrane bending

We next asked how the membrane remodeling ability of crowded disordered domains compares with that of BAR scaffolds. To make this comparison, we created a chimeric protein that places the two mechanisms in direct competition within the same molecule. Specifically, we fused the inverted BAR (I-BAR) domain of human IRSp53 to the bulky, C-terminal disordered domain of rat AP180 (569 disordered amino acids) to form I-BAR-AP180 CTD (Fig. 5A). While the I-BAR domain is known to generate inverted membrane curvature (Mattila et al., 2007; Saarikangas et al., 2009), the attached disordered domains should generate steric pressure that will tend to bend the membrane in the opposite direction. Exposing GUVs to the I-BAR domain alone drove inverted membrane tubulation, as expected (Fig. 5B, left and Movie S6). In contrast, the I-BAR-AP180 CTD chimera drove neither inward nor outward tubulation. Instead, the protein caused the GUV membrane to fluctuate dynamically through non-spherical shapes in which regions of gentle membrane curvature extending both inward and outward were apparent, (Fig. 5B, right and Movie S7). These “frustrated” fluctuations demonstrate that the disordered domain effectively neutralized the ability of the I-BAR domain to drive inward membrane bending. This result suggests that crowding among disordered domains and scaffolding by BAR domains make comparable contributions to membrane remodeling.

**Figure 5.**
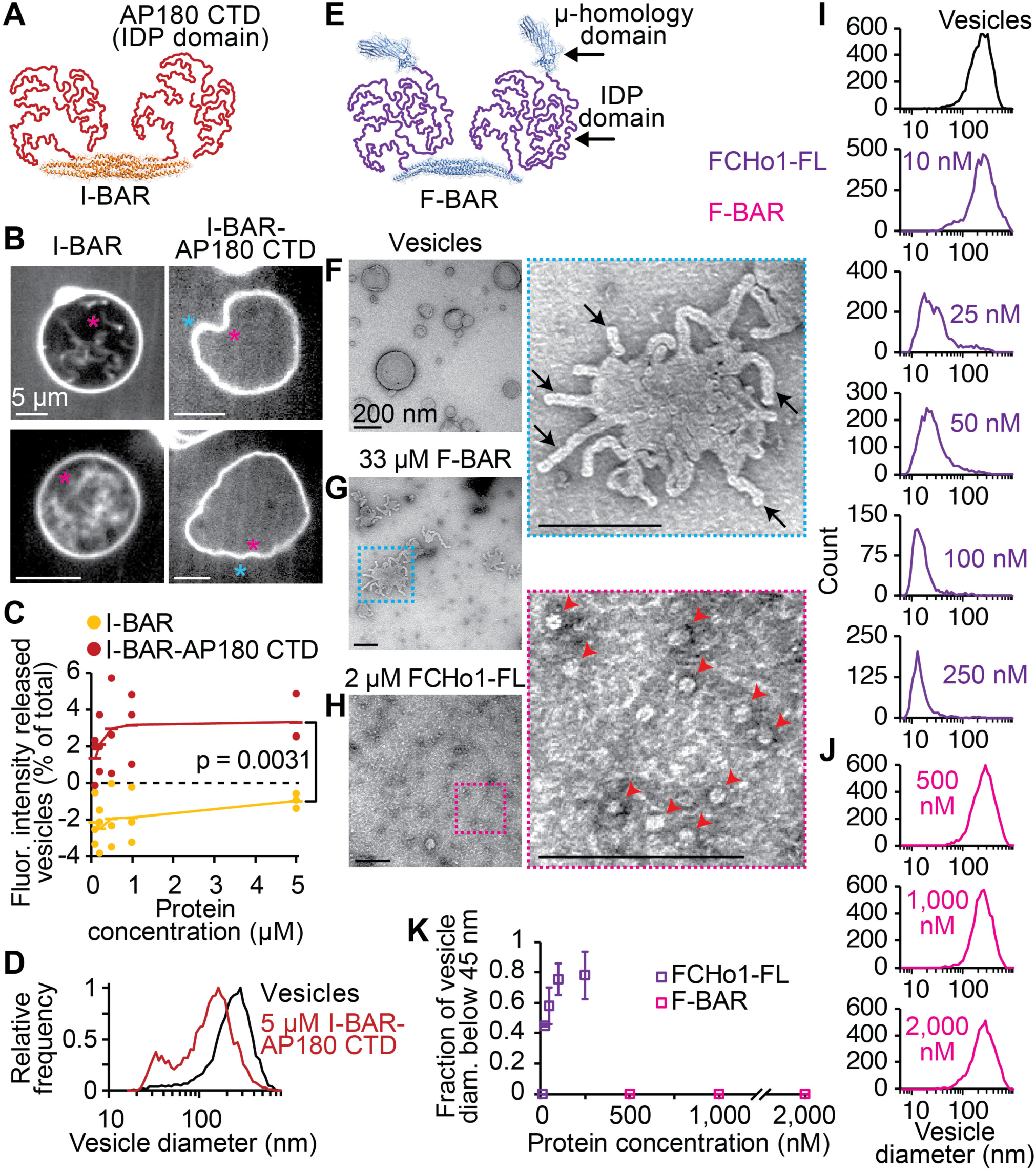
Disordered domain crowding opposes inverted membrane bending by I-BARs and promotes membrane fission by F-BARs. GUV membrane composition: 79.5 mol% DOPC, 5 mol% PtdIns(4,5)P2, 15 mol% DOPS, 0.5 mol% Oregon Green 488-DHPE. SUPER template membrane composition: 79 mol% DOPC, 5 mol% PtdIns(4,5)P_2_, 15 mol% DOPS, 1 mol% Texas Red-DHPE. Tethered vesicle membrane composition: 76 mol% DOPC, 5 mol% PtdIns(4,5)P2, 15 mol% DOPS, 2 mol% Oregon Green 488-DHPE, 2 mol% DP-EG10-biotin.

Membrane release experiments using SUPER templates showed that exposure to the I-BAR domain (100 nM - 5 μM) decreased the amount of membrane shedding to levels lower than the amount of non-specific shedding that occurred in protein-free buffer (Fig. 5C, negative values). This result suggests that assembly of the I-BAR scaffold stabilized the membrane against shedding. In contrast, I-BAR-AP180 CTD drove positive membrane release at all concentrations (Fig. 5C), demonstrating that the disordered domain of AP180 is capable of disrupting the membrane-stabilizing effect of I-BAR. Interestingly, tethered vesicle fission experiments revealed that the highest concentration of I-BAR-AP180 CTD that we tested, 5 μM, drove membrane fission, generating a population of vesicles centered near 30 nm diameter (Fig. 5D). This result demonstrates that, under appropriate conditions, steric pressure among crowded disordered domains is sufficient to overcome the structure-based curvature preference of the I-BAR scaffold. Importantly, while wild-type IRSp53 does not naturally contain a large disordered domain, the I-BAR domain-containing proteins MIM and ABBA do contain regions of substantial disorder (approximately 475 amino acids in MIM) (Lee et al., 2007). Therefore, our observations raise the question of whether the presumed role of these proteins in driving cellular membrane protrusions can be justified on the basis of in vitro studies of their isolated I-BAR domains (Mattila et al., 2007; Saarikangas et al., 2009).

### An F-BAR scaffold drives fission by locally crowding bulky disordered domains

If BAR scaffolds drive membrane fission by concentrating large disordered domains at membrane surfaces, then the ability of Amph-FL to drive fission (Figs. 1 and 2) should extend to other proteins that contain both BAR domains and substantial regions of intrinsic disorder. Interestingly, many proteins that contain the modestly-curved F-BAR domain also have this architecture (Roberts-Galbraith and Gould, 2010), including the endocytic proteins FCHo1/2 (Henne et al., 2010) and its yeast homolog Syp1 (Reider et al., 2009), the srGAP proteins involved in neuronal development (Wuertenberger and Groemping, 2015), and the cytokinesis proteins Cdc15 (Roberts-Galbraith et al., 2010) and Imp2 (McDonald et al., 2016) in S. *pombe* and Hof1 (Meitinger et al., 2011) in S. *cerevisiae*. To test this idea, we examined FCHo1 (C. *elegans)*, which consists of an N-terminal F-BAR domain followed by an intrinsically disordered domain of 412 amino acids and a C-terminal μ-homology domain (Henne et al., 2010; Ma et al., 2016; Umasankar et al., 2014; Wang et al., 2016) (Fig. 5E). Negative stain TEM revealed that exposure of vesicles with an initial average diameter of 200 nm to the F-BAR domain of FCHo1 drove formation of lipid tubules with an average diameter of 21±2 nm s.d. (Fig. 5F,G and S5A,B), in agreement with previous findings (Henne et al., 2010; Henne et al., 2007). In contrast, full-length FCHo1 (FCHo1-FL) did not generate lipid tubules, but instead divided the 200 nm diameter vesicles into a population of highly curved vesicles with average diameter 17±7 nm s.d. (Fig. 5H and S5C,D).

Similarly, tethered vesicle fission experiments revealed that FCHo1-FL drove potent membrane fission over the concentration range of 10-250 nM (Fig. 5I,K), a comparable range to Amph-FL (50-150 nM, Fig. 2D,F), with a mean diameter of 19±1 nm s.e.m. at 250 nM of FCHo1-FL. However, the F-BAR domain alone did not drive fission in these experiments, even at substantially higher concentrations up to 2 μM (Fig. 5J,K). Interestingly, in both tethered vesicle and TEM experiments, the fission products produced by FCHo1-FL had a peak diameter of slightly less than 20 nm, with some vesicles as small as 10 nm. These small diameters suggest that FCHo1-FL may stabilize the formation of membrane micelles, which are similar in morphology to the hemifusion intermediates that form during membrane fission (Campelo et al., 2012; Frolov et al., 2015). Notably, vesicles in these experiments were composed primarily of the lipid DOPC, which has a lower bending rigidity than physiological membranes (Dimova, 2014). In contrast, when we increased bilayer rigidity by replacing DOPC with the substantially more rigid, fully-saturated lipid DPPC (Dimova, 2014; Lee et al., 2001), the average diameter of fission products increased to approximately 50 nm at 50 nM of FCHo1-FL, consistent with bilayer vesicles (Fig. S5E-G). Collectively, these studies show that, despite their gentle curvature, F-BAR scaffolds are also capable of collaborating with disordered domain crowding to drive efficient membrane fission, producing highly curved vesicles. These findings suggest that the ability of BAR domains to assemble into scaffolds that concentrate disordered domains, regardless of their intrinsic structural curvature, is the key requirement for membrane fission.

Membrane composition for vesicles in TEM: 80 mol% DOPC, 5 mol% PtdIns(4,5)P_2_, 15 mol% DOPS. (**A**) Schematic of I-BAR-AP180 CTD chimera dimer. IRSp53 I-BAR domain: PDB 1Y2O. (**B**) Two representative confocal micrographs of GUVs after exposure to I-BAR or I-BAR-AP180 CTD. Asterisks indicate direction of membrane bending (magenta: inward; cyan: outward). Fluorescence signal comes from Atto 594-labeled protein. (**C**) SUPER template membrane release comparing I-BAR and I-BAR-AP180 CTD. Dots indicate data and lines indicate mean, n = 3 independent experiments. P-value: one-tailed, unpaired Student’s t-test. (**D**) Tethered vesicle fission experiments reveal that 5 μM I-BAR-AP180 CTD generates highly curved fission vesicles. (**E**) Schematic of full-length FCHo1 (FCHo1-FL) dimer. F-BAR domain: PDB 2V0O. μ-homology domain: PDB 5JP2, chain A. (**F-H**) Negative stain TEM micrographs of 200 nm extruded vesicles (**F**) before exposure to protein, (**G**) after exposure to 33 μM F-BAR, and (**H**) after exposure to 2 μM FCHo1-FL. Dashed boxes indicate zoomed regions to the right. Black arrows indicate membrane tubules, red arrowheads indicate fission vesicles. (**I**) Tethered vesicle fission experiments reveal FCHo1-FL generates highly curved fission products over the concentration range of 10-250 nM. (**J**) F-BAR does not drive fission in tethered vesicle fission experiments, even at concentrations up to 2,000 nM. (**K**) Plot of ratio of vesicle distribution area below 45 nm to the total distribution area (compare to Fig. 2F). Markers in represent mean ± first s.d., n = 3 independent experiments. Scale bars in (B): 5 μm. Scale bars in (F-H), including zoomed regions: 200 nm. See also Fig. S5 and Movies S6 and S7.

## Discussion

Here we report that membrane scaffolding by BAR domains works synergistically with steric pressure among bulky disordered domains to drive membrane fission. By highlighting the ability of BAR scaffolds to locally concentrate disordered domains, this work helps to explain how steric pressure can be generated and locally sustained at membrane surfaces. Further, our findings are in contrast with the established view that BAR scaffolds prevent fission by stabilizing membrane tubes (Boucrot et al., 2012). Instead, our work suggests that BAR proteins that contain substantial disordered regions are more likely to be drivers of membrane fission. In support of this idea, previous work showed that the yeast amphiphysins Rvs161/167 are essential for membrane fission, and deletion of these proteins leads to a defect in the entry of clathrin-coated pits into cells (Kaksonen et al., 2005; Kishimoto et al., 2011). Depletion of amphiphysin by RNA interference also dramatically inhibits clathrin-mediated endocytosis in mammalian cells (Meinecke et al., 2013), although this endocytic defect has been attributed to a reduction in dynamin recruitment.

Our finding that modestly-curved F-BAR domains can also collaborate with bulky disordered domains to drive potent fission is surprising, as it suggests that proteins such as FCHo1/2 may participate in membrane shaping throughout the maturation of clathrin-coated pits, and may even help drive the final fission event. This hypothesis is in contrast to the idea that FCHo1/2 are primarily involved in the initiation of clathrin-coated pits (Henne et al., 2010; Ma et al., 2016). However, previous work showed that FCHo2 is present throughout the lifetime of clathrin-coated pits (Taylor et al., 2011), supporting the possible role of FCHo2 in membrane shaping. Moreover, many F-BAR proteins involved in other cellular pathways such as cytokinesis also contain large regions of intrinsic disorder (McDonald et al., 2016; Meitinger et al., 2011; Roberts-Galbraith et al., 2010). As such, our findings raise the question of whether F-BAR scaffolds facilitate membrane fission in a variety of contexts beyond membrane traffic.

More broadly, our work raises the possibility that protein assemblies beyond BAR domains may serve as scaffolds for crowding bulky disordered domains in order to ensure efficient membrane fission. One potential example is dynamin, a scaffold-forming GTPase involved in fission of clathrin-coated pits (Antonny et al., 2016). While dynamin itself does not contain substantial regions of disorder, it assembles with proteins that contain such domains, including amphiphysin and SNX9 (Daumke et al., 2014). Once recruited by dynamin, these proteins may generate significant steric pressure at membrane necks. In line with this thinking, previous in vitro studies found that amphiphysin acts synergistically with dynamin to enhance membrane fission (Meinecke et al., 2013; Neumann and Schmid, 2013). Moreover, the yeast dynamin homolog, Vps1, is dispensable for fission, but is necessary for proper localization and accumulation of amphiphysin at clathrin-coated pits (Kaksonen et al., 2005; Smaczynska-de Rooij et al., 2010). A function of Vps1 could therefore be to organize and concentrate bulky disordered domains at membrane necks to promote fission.

Recent work has revealed that BAR domains also promote membrane fission by acting as a diffusion barrier to lipids (Renard et al., 2015; Simunovic et al., 2017). This “friction-driven scission” mechanism may be responsible for fission in a clathrin-independent endocytic pathway (Renard et al., 2015; Simunovic et al., 2017). While friction-driven scission and disordered domain crowding are distinct mechanisms, they are not mutually exclusive and may therefore work cooperatively to drive fission of endocytic structures. Future work is needed to better understand the potential collaboration between these BAR domain-mediated fission mechanisms.

Our work reveals a synergistic relationship between structured protein assemblies and disordered pressure generators, which can be harnessed to drive membrane fission. It is increasingly recognized that structural disorder is prevalent in membrane trafficking, and that disordered domains are often coupled to structured domains within the same protein molecules (Pietrosemoli et al., 2013). While previous work has focused primarily on structure-function relationships revealed by studying individual protein domains, these findings highlight the importance of examining the collective contributions from both structure and disorder to understand how proteins shape membranes in diverse cellular contexts.

## Materials and Methods

### Chemical reagents

MOPS (3-(N-morpholino)propanesulphonic acid), HEPES (4-(2-hydroxyethyl)-1-piperazineethanesulphonic acid), Tris hydrochloride, NaCl, DTT (dithiothreitol), IPTG (isopropyl-β-D-thiogalactopyranoside), β-mercaptoethanol, thrombin protease, and Triton X-100 were purchased from Fisher Scientific. EDTA (ethylenediaminetetraacetic acid), EGTA (ethylene glycol tetraacetic acid), TCEP (tris(2-carboxyethyl)phosphine hydrochloride), PMSF (phenylmethanesulfonyl fluoride), EDTA-free protease inhibitor tablets, Thrombin CleanCleave Kit, PLL (poly-L-lysine), Atto 488 NHS-ester, and Atto 594 NHS-ester were purchased from Sigma-Aldrich. HRV-3C (human rhinovirus-3C) protease, neutravidin, Oregon Green 488-DHPE (Oregon Green 488 1,2-dihexadecanoyl-*sn*-glycero-3-phosphoethanolamine), and Texas Red-DHPE were purchased from Thermo Fisher Scientific. mPEG-SVA (mPEG-succinimidyl valerate), biotin-PEG-SVA, mPEG-silane, and biotin-PEG-silane (all PEGs were molecular weight 5,000 Da) were purchased from Laysan Bio (Arab, AL, USA). DP-EG10-biotin (dipalmitoyl-decaethylene glycol-biotin) was generously provided by Dr. Darryl Sasaki of Sandia National Laboratories, Livermore, CA (Momin et al., 2015). All other lipids were purchased from Avanti Polar Lipids (Alabaster, AL, USA), including: PtdIns(4,5)P_2_ (L-α- phosphatidylinositol-4,5-bisphosphate, from porcine brain), DOGS-NTA-Ni (1,2-dioleoyl-sn-glycero-3-[(N-(5-amino-1-carboxypentyl)iminodiacetic acid)succinyl], nickel salt), DOPC (1,2-dioleoyl-sn-glycero-3-phosphocholine), and DOPS (1,2-dioleoyl-sn-glycero-3-phospho-L-serine, sodium salt). The lipid compositions for all experiments are listed in the figure captions.

### Plasmids

The pGex6P bacterial expression vector containing full-length human amphiphysin (residues 2-695) was generously provided by the Baumgart Lab (University of Pennsylvania). The N-BAR domain of human amphiphysin (residues 2-242) was cloned into the pGex4T2 bacterial expression vector using BamHI and EcoRI restriction sites. The C-terminal domain of human amphiphysin lacking the SH3 domain (Amph CTD DSH3, residues 240-622) with N-terminal GST and 6his tags was cloned using a previously-generated plasmid template, GST-6his-AP180 CTD in pGex4T2 (Busch et al., 2015). AP180 CTD was excised from the template using SalI and XhoI restriction sites and the Amph CTD ∆SH3 insert was ligated in using the same SalI and XhoI sites. The N-BAR domain of human amphiphysin fused to the C-terminal domain of rat epsin1 (N-BAR-epsin CTD, residues 144-575 of rat epsin1) was cloned by first ligating in the N-BAR domain of human amphiphysin (residues 2-242) into pGex4T2 using BamHI and EcoRI restriction sites. Epsin CTD was then ligated in frame with N-BAR using SalI and NotI restriction sites. The I-BAR domain of human IRSp53 (residues 1-250) was cloned by using site directed mutagenesis to introduce a stop codon at residue 251 in the pGex6P2 plasmid containing full-length IRSp53. The I-BAR domain of human IRSp53 fused to the C-terminal domain of rat AP180 (I-BAR-AP180 CTD, residues 328-896 of rat AP180) was cloned by first ligating the I-BAR domain of human IRSp53 (residues 1-250) into pGex4T2 using BamHI and EcoRI restriction sites. AP180 CTD was then ligated in frame with I-BAR using SalI and XhoI restriction sites. The pGex6P1 vector containing full-length *C. elegans* FCHo1 (residues 1-968) was generously provided by the Audhya Lab (University of Wisconsin - Madison) (Wang et al., 2016). The F-BAR domain of *C. elegans* FCHo1 (residues 1-276) was cloned into the pGex4T2 vector using BamHI and EcoRI restriction sites.

The pCAGEN mammalian expression vector containing the N-BAR domain of human amphiphysin (residues 1-256), tagged at the C-terminus with mCherry, was a gift from Tobias Meyer (Addgene plasmid # 85130). Full-length human amphiphysin (residues 1-695) was cloned into the pCAGEN vector, in frame with mCherry at the C-terminus, by first excising the N-BAR domain from the template using EcoRI and AgeI restriction sites, and then ligating in full-length amphiphysin using the same EcoRI and AgeI restriction sites. The N-BAR domain of human amphiphysin fused to the C-terminal domain of mouse neurofilament-M (N-BAR-NfM CTD, residues 411-848 of mouse neurofilament-M) was cloned by ligating neurofilament-M CTD into the existing N-BAR-mCherry pCAGEN template, between N-BAR and mCherry, using a single AgeI restriction site. The resulting plasmid contained a GPV linker between N-BAR and neurofilament-M CTD and a GPVAT linker between neurofilament-M CTD and mCherry. All plasmids were confirmed by DNA sequencing.

### Protein purification

All proteins were expressed as N-terminal glutathione-S-transferase (GST) fusion constructs in BL21 *E. coli* cells following induction with 1 mM IPTG. Full-length amphiphysin, N-BAR, Amph CTD ΔSH3, I-BAR, I-BAR-AP180 CTD and F-BAR were induced at 30 °C for 6-8 h. N-BAR-epsin CTD was induced at 16 °C for 20 h. Full-length FCHo1 was induced at 12 °C for 24 h. Cells were harvested and bacteria were lysed using lysis buffer and probe sonication. For full-length FCHo1, lysis buffer was: 100 mM sodium phosphate pH 8.0, 5 mM EDTA, 5 mM DTT, 10% glycerol, 1 mM PMSF, 1% Triton X-100, 1x Roche protease inhibitor cocktail. For all other proteins, lysis buffer was: 500 mM Tris-HCl pH 8.0, 5 mM EDTA, 10 mM β-mercaptoethanol or 5 mM TCEP, 5% glycerol, 1 mM PMSF, 1% Triton X-100, 1x Roche or Pierce protease inhibitor cocktail. Proteins were purified from bacterial extracts by incubating with glutathione resin, followed by extensive washing (at least 10x column volumes). Full-length amphiphysin, N-BAR, Amph CTD ΔSH3, and F-BAR were cleaved directly from the resin using soluble HRV-3C (Thermo Fisher) or thrombin (GE Healthcare Life Sciences) proteases overnight at 4 °C with rocking. HRV-3C, which contained a GST tag, was removed by passage through a glutathione agarose column. Thrombin was removed with *p*-aminobenzamidine-agarose resin (Sigma-Aldrich). N-BAR-epsin CTD, I-BAR, and I-BAR-AP180 CTD were eluted with 15 mM reduced glutathione in 500 mM Tris-HCl pH 8.0, 5 mM EDTA, 10 mM β-mercaptoethanol or 5 mM TCEP, 5% glycerol, 1 mM PMSF buffer. Full-length FCHo1 was eluted with 15 mM reduced glutathione in 100 mM sodium phosphate pH 8.0, 5 mM EDTA, 5 mM DTT, 10% glycerol, 1 mM PMSF buffer. The proteins were concentrated with EMD Millipore Amicon centrifugal filter units, desalted with Zeba Spin Desalting Columns (Fisher), and then incubated with either Thrombin CleanCleave Kit (Sigma-Aldrich), soluble HRV-3C, or soluble thrombin overnight at 4 °C with rocking. Cleaved GST was removed by passage through a glutathione agarose column. I-BAR-AP180 CTD and N-BAR-epsin CTD were further purified by gel filtration chromatography using a Superose 6 column equilibrated with 20 mM Tris-HCl pH 8.0, 150 mM NaCl, 1 mM EDTA, 5 mM EGTA, 1 mM PMSF, 5 mM DTT. All proteins were stored as small aliquots or liquid nitrogen pellets at −80 °C.

### Protein labeling

Proteins were labeled using amine-reactive, NHS ester-functionalized dyes (Atto-Tec) in 25 mM HEPES pH 7.35, 150 mM NaCl, 5 mM TCEP buffer. The concentration of dye was adjusted experimentally to obtain the desired labeling ratio of 0.5-1 dye molecules per protein, typically 2-5 times molar excess of dye. Reactions were performed for 20-30 min at room temperature and labeled protein was separated from unconjugated dye using Princeton CentriSpin-20 size exclusion spin columns (Princeton Separations).

### Transmission electron microscopy

Vesicles for electron microscopy were composed of 5 mol% PtdIns(4,5)P_2_, 15 mol% DOPS, and 80 mol% DOPC. Dried lipid films were hydrated in 20 mM MOPS pH 7.35, 150 mM NaCl, 0.5 mM EGTA and EDTA buffer and extruded though a 200 nm pore filter (Whatman). Proteins were diluted to the indicated concentrations in the same MOPS buffer with 5 mM TCEP and incubated with vesicles at 37 °C for 30 min (Amph-FL and N-BAR) or 60 min (FCHo1-FL and F-BAR). The vesicle concentration was 1 mM in experiments with Amph-FL, FCHo1-FL, and F-BAR, and 0.1 mM in experiments with N-BAR and in protein-free controls. 5 μL of the mixture was placed onto a glow-discharged, 300 square mesh, carbon-coated grid and stained with 2% uranyl acetate (Electron Microscopy Sciences; Hatfield, PA, USA). Images were collected on a Tecnai Spirit BioTwin T12 electron microscope (Tecnai; Hillsboro, OR, USA). Vesicle and tubule diameters were measured using ImageJ software.

### Fluorescence microscopy

A spinning disc confocal microscope (Zeiss Axio Observer Z1 with Yokagawa CSU-X1M) was used to image GUVs and tethered vesicles. Laser wavelengths of 488 and 561 nm were used for excitation. Emission filters were centered at 525 nm with a 50 nm width, and 629 nm with a 62 nm width. A triple-pass dichroic mirror was used: 405/488/561 nm. The microscope objective was a Plan-Apochromat 100x, 1.4 numerical aperture oil immersion objective. Images were collected on a cooled (−70 °C) EMCCD iXon3 897 camera (Andor Technology; Belfast, UK).

### Giant unilamellar vesicle preparation

Giant unilamellar vesicles (GUVs) were prepared according to published protocols (Angelova and Dimitrov, 1986). A mixture of 5 mol% PtdIns(4,5)P_2_, 15 mol% DOPS, 0.5 mol% Oregon Green 488-DHPE, and 79.5 mol% DOPC was dried into a film on an indium-tin-oxide coated glass slide. Lipids were further dried under vacuum overnight. Electroformation was performed at 55 °C in 350 mOsm sucrose solution also containing 0.5 mM EGTA and EDTA to prevent clustering of PtdIns(4,5)P_2_. Vesicles were mixed with protein solution at the specified concentration in 20 mM MOPS pH 7.35, 150 mM NaCl, 0.5 mM EGTA and EDTA, 5 mM TCEP buffer. Prior to mixing, the osmolarity of the GUV solution and experiment buffer was measured using a vapor pressure osmometer (Wescor).

### SUPER template preparation

SUPER templates were prepared according to the protocol of Neumann et al (Neumann et al., 2013). A lipid mixture of 5 mol% PtdIns(4,5)P_2_, 15 mol% DOPS, 1 mol% Texas Red DHPE, and 79 mol% DOPC was mixed in a clean glass test tube, the solvent was evaporated, and the lipid film was further dried under vacuum. The lipid film was hydrated in Milli-Q water, subjected to three freeze-thaw cycles in liquid nitrogen, and extruded through a 100 nm pore filter (Whatman). SUPER templates were made by creating a 100 μL mixture consisting of 200 μM liposomes, 1 M NaCl, and 5 × 10^6^ of 2.5 μm m-type silica beads (Corpuscular; Cold Spring, NY, USA) in a low-adhesion microcentrifuge tube. The mixture was incubated for 30 min at room temperature and gently agitated periodically. The mixture was washed by adding 1 mL Milli-Q water, gently mixing, and spinning at 300 g for 2 min in a swinging bucket rotor to pellet the SUPER templates. 1 mL of supernatant was removed, SUPER templates were resuspended in the remaining 100 μL, and washing was repeated a total of four times. SUPER templates were kept on ice and used within 4 h.

### Measurement of SUPER template membrane release

SUPER template membrane shedding experiments were performed according to the protocol of Neumann et al (Neumann et al., 2013). 10 μL of SUPER templates were gently pipetted into the top of a 90 μL solution of protein at specified concentrations in 20 mM MOPS pH 7.35, 150 mM NaCl, 0.5 mM EGTA and EDTA, 5 mM TCEP buffer. SUPER templates were allowed to slowly settle for 30 min at room temperature without disturbing. SUPER templates containing unreleased membrane were then sedimented by gentle centrifugation at 300 g for 2 min in a swinging bucket rotor. 75 μL of supernatant containing released membrane was collected and mixed in a 96-well plate with Triton X-100 at a final concentration of 0.1% and volume of 100 μL. In order to measure the total fluorescence of SUPER template membrane, a detergent control consisting of SUPER templates added directly to 0.1% Triton X-100, which solubilized all SUPER template membrane, was run. The fluorescence intensity of released membrane was measured in a plate reader using 590 nm excitation light and an emission filter centered at 620 nm. After subtracting the fluorescence of 0.1% Triton X-100 in buffer alone from all measurements, membrane release was calculated by dividing the fluorescence intensity after protein exposure by the fluorescence intensity of the detergent control. The background level of membrane release in the absence of protein was also measured by incubating SUPER templates in buffer alone. This buffer control was subtracted from all measurements as background.

### Fluorescence correlation spectroscopy

Imaging wells for fluorescence correlation spectroscopy (FCS) utilized supported lipid bilayers (SLBs) to passivate the glass surface and prevent protein adsorption. Briefly, a well was created with a silicone gasket on an ultraclean coverslip, and a solution of sonicated DOPC vesicles at 1 mM lipid was added. The SLB was formed for 10 min and thoroughly washed in experiment buffer of 50 mM Tris pH 8.0, 10 mM CaCl_2_, 150 mm NaCl, 15 mM EGTA, 5 mM EDTA, 5 mM TCEP. Atto 488-labeled proteins were diluted in experiment buffer and added to the imaging well such that the concentration of Atto 488 dye was approximately 1 nM. FCS measurements were acquired on a custom-built time-correlated single photon counting confocal microscope using a 486 nm picosecond pulsed diode laser. The laser was focused in solution approximately 3 μm above the bilayer passivation surface, and fluorescence signal was collected as proteins diffused through the focused laser volume. The signal was split onto separate GaAsP photomultiplier tubes (Hamamatsu) for cross-correlation using Becker and Hickl software. FCS traces were collected for 120 s. The number of FCS traces acquired for Amph CTD ∆SH3, AP180 CTD, and transferrin were: 10, 5, and 3, respectively. Each FCS trace was fit with the 2D autocorrelation function:

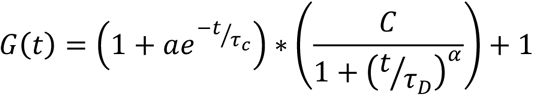

Where *C* is *1/N_p_*, *N_p_* is the number of labeled proteins in the focused laser volume, *τ_D_* is the diffusion time, and *α* is the anomalous diffusion coefficient. *a* and *τ_c_*, which correct for short time processes such as intersystem crossing, were held constant in the fitting as 0.05 and 5 μs, respectively (Houser et al., 2016). Fitting was performed in Wolfram Mathematica 11 software. *α* values were between 0.90 and 0.93 for all fits, demonstrating that a substantial correction for anomalous diffusion was not needed.

Fig. S2G-I shows representative FCS traces and fits for Amph CTD ∆SH3, AP180 CTD, and transferrin, with mean values of *τ_D_* ± first s.d. reported next to each trace. The hydrodynamic radius, *R_H_*, of each protein is also reported next to each trace. AP180 CTD was used as a calibration standard to compute *R_H_* of Amph CTD ∆SH3, as *τ_D_* is directly proportional to *R_H_*. AP180 CTD is a disordered protein with a radius that has been well-characterized (Busch et al., 2015; Kalthoff et al., 2002), making it an appropriate calibration standard. This calibration approach also yielded a radius for transferrin that was consistent with its expected radius (Hall et al., 2002), confirming the validity of our approach. The radius of Amph CTD ∆SH3 was taken as 5 nm in estimates of membrane coverage by proteins in Fig. 3D, S2A-F, and S2K-M. This value was chosen based on previous studies which found that the radius of gyration of a disordered protein is approximately 1.2-fold greater than the hydrodynamic radius (Hofmann et al., 2012; Sherman and Haran, 2006).

Fig. S2J shows the relative diffusion time, *τ_D_*, of Amph CTD ∆SH3 in 20 mM MOPS pH 7.35 in varying concentrations of NaCl. Data are plotted as the relative diffusion time compared to the diffusion time at 150 mM NaCl. Diffusion times were corrected for changes in solution viscosity with changing NaCl concentration (Zhang and Han, 1996). Because *τ_D_* is directly proportional to *R_H_*, the increase in *τ_D_* at 10 mM NaCl indicates an increase in *R_H_*, likely owing to reduced charge screening which expands and extends the disordered protein (Srinivasan et al., 2014). Similarly, the decrease in *τ_D_* at 1 M NaCl indicates a decrease in *R_H_*, owing to enhanced charge screening which compacts the disordered protein.

### Passivating glass coverslips with PEG and PEG-biotin for tethering vesicles

Glass coverslips were passivated by either directly conjugating PEG-silane and biotin-PEG-silane to the glass, or by coating the glass with a layer of poly-L-lysine (PLL) conjugated to PEG and biotin-PEG. For the direct silane conjugation, a 0.67% solution of PEG-silane was prepared in anhydrous isopropanol. Biotin-PEG-silane comprised 5% of the total amount of PEG-silane in the solution. The mixture was held in a bath sonicator for 10-15 min to dissolve the PEG. Acetic acid was added to a concentration of 1%, and 50 μL of the reactive mixture was dropped onto a dry, ultraclean coverslip. Another dry, ultraclean coverslip was sandwiched on top, and the slides were incubated at 70 °C for 30-60 min. The slides were separated, washed in ultrapure water, and stored dry for later use. Imaging wells were made by placing silicone gaskets onto the glass and hydrating in 20 mM MOPS pH 7.35, 150 mM NaCl buffer. Neutravidin was added to the well at a final concentration of 0.2 mg mL^-1^, incubated for 10 min, and the well was washed repeatedly with MOPS buffer before adding vesicles.

The biotinylated PLL-PEG was made according to a previous protocol (Ruiz-Taylor et al., 2001). Briefly, amine-reactive PEG-SVA (succinimidyl valerate) and biotin-PEG-SVA was added to a 40 mg mL^-1^ mixture of PLL in 50 mM sodium tetraborate pH 8.5 at a molar ratio of one PEG per five lysine subunits. PEG-biotin comprised 2% of the total PEG amount. The mixture was stirred continuously for 6 h at room temperature and buffer exchanged into PBS using Centri-Spin size exclusion columns (Princeton Separations). Imaging wells were made by placing silicone gaskets onto ultraclean coverslips. Wells were coated for 20-30 min with biotinylated PLL-PEG diluted tenfold in 20 mM MOPS pH 7.35, 150 mM NaCl buffer. After coating, the well was washed repeatedly with MOPS buffer to wash out excess PLL-PEG. Neutravidin was added to the well following the same process as for PEG-silane slides.

### Determination of vesicle diameter from measurements of tethered vesicle brightness

Vesicle diameter distributions were measured using an assay developed by the Stamou group (Hatzakis et al., 2009; Kunding et al., 2008; Stamou et al., 2003). Vesicles in experiments with Amph-FL, N-BAR, N-BAR-epsin CTD, I-BAR-AP180 CTD, FCHo1-FL, and F-BAR were composed of 76 mol% DOPC, 15 mol% DOPS, 5 mol% PtdIns(4,5)P_2_, 2 mol% DP-EG10-biotin, and 2 mol% Oregon Green 488-DHPE. Vesicles in experiments with Amph CTD ΔSH3 were composed of a similar lipid mixture, with the exception that DOPS and PtdIns(4,5)P_2_ were replaced with 20 mol% DOGS-NTA-Ni. Experiments with Amph-FL and N-BAR on highly charged membranes (Fig. S3) used vesicles composed of 68 mol% DOPS, 23 mol% DOPE, 5 mol% cholesterol, 2 mol% DP-EG10-biotin, and 2 mol% Oregon Green 488-DHPE, similar to Mim et al, Cell 2012 (Mim et al., 2012). Dried lipid films were hydrated in 20 mM MOPS pH 7.35, 150 mM NaCl buffer (0.5 mM EGTA and EDTA was included in experiments with PtdIns(4,5)P_2_) and extruded to 200 nm.

Fission experiments were performed by mixing vesicles at a concentration of 10 μM with unlabeled protein at specified concentrations in the above MOPS buffer with 5 mM TCEP. The mixture was incubated at 37 °C for either 30 min (Amph-FL, N-BAR, Amph CTD ΔSH3, and N-BAR-epsin CTD) or 60 min (I-BAR-AP180 CTD, FCHo1-FL, and F-BAR). During the incubation period, imaging wells were prepared as described above. After incubation, the mixtures were added to the wells and vesicles were allowed to tether for 10 min before washing repeatedly to remove untethered vesicles. Multiple spinning disc confocal z-stacks of tethered vesicles were acquired with a z-step of 0.1 μm. The same laser power and camera gain settings were used for all experiments.

All images in the z-stacks were cropped to the center 171×171 pixels (center 1/9^th^), and the frame with the greatest mean brightness was selected as the best focus image for analysis. Fluorescence amplitudes of diffraction-limited puncta were obtained using cmeAnalysis particle detection software (Aguet et al., 2013). Individual vesicles were detected by fitting two-dimensional Gaussian profiles to each puncta. The standard deviation of the Gaussian was determined from the point spread function of our microscope. The brightness values of detected puncta were reported as valid if they were diffraction-limited and had amplitudes significantly above their local fluorescence background. To further ensure that puncta were well above the noise threshold, we only accepted puncta that persisted at the same location through five consecutive imaging frames.

To convert fluorescence brightness values to vesicle diameters, we computed a scaling factor that centered the mean of the vesicle brightness distribution of a high-curvature, sonicated vesicle sample to the average diameter of the same vesicles obtained from dynamic light scattering. This scaling factor was then used to scale the vesicle brightness distributions after protein exposure to distributions of vesicle diameter.

### Determination of membrane coverage by proteins from measurements of vesicle and protein brightness

Vesicles in experiments with full-length amphiphysin and N-BAR were composed of 76 mol% DOPC, 15 mol% DOPS, 5 mol% PtdIns(4,5)P_2_, 2 mol% DP-EG10-biotin, and 2 mol% Oregon Green 488-DHPE. DOPS and PtdIns(4,5)P_2_ were replaced with 20 mol% DOGS-NTA-Ni in experiments with Amph CTD ∆SH3. Dried lipid films were hydrated in 20 mM MOPS pH 7.35, 150 mM NaCl buffer (0.5 mM EGTA and EDTA was included in experiments with PtdIns(4,5)P_2_) and extruded to 30 or 200 nm. Imaging wells were prepared as described above. Vesicles were diluted to 5 μM in the wells and allowed to tether for 10 min. Untethered vesicles were removed by thorough washing with MOPS buffer. After tethering, Atto 594-labeled protein was added to the specified concentration, and multiple spinning disc confocal z-stacks of lipid and protein fluorescence were acquired, with a z-step of 0.1 μm. Images were collected after approximately 15 min incubation of protein with vesicles. The same laser power and camera gain settings were used for all experiments. Fig. S2A shows images of tethered vesicles with 10 and 25 nM Amph-FL-Atto 594, demonstrating increased protein brightness (and therefore membrane coverage) with increasing protein concentration.

Images were cropped and individual vesicle puncta were detected using cmeAnalysis software (Aguet et al., 2013), following a similar approach described in the previous section. Here we only accepted puncta that persisted at the same location through three consecutive imaging frames. The algorithm also searched for fluorescent puncta in the protein channel using the centroids of the detected fluorescent puncta in the master lipid channel. The search region in the protein channel was three times the standard deviation of the Gaussian fit to the point spread function of our microscope. We estimated vesicle diameters from lipid fluorescence brightnesses by calibrating against dynamic light scattering, as described in the previous section. We estimated the number of bound proteins on each vesicle by comparing brightness values in the protein channel to the brightness of a single molecule of Atto 594-labeled protein. Images of single molecules of Atto 594-labeled proteins were obtained by adding a dilute concentration of protein to an imaging well on an ultraclean coverslip, and imaging single proteins adhered to the coverslip surface in a similar manner as described for the tethered vesicles. A linear correction for camera exposure time was applied to the single molecule brightness, as longer exposure times were required to image single molecules compared to membrane-bound protein. Fig. S2B shows a plot of the raw protein intensity values as a function of vesicle intensity for 10 and 25 nM Amph-FL. The 25 nM data shows a higher slope than 10 nM, indicating greater membrane coverage. Fig. S2C shows this same data after processing, plotted as the number of membrane-bound proteins as a function of vesicle diameter.

Membrane coverage by proteins was estimated for each vesicle by dividing the area occupied by membrane-bound proteins by the corresponding vesicle surface area. The projected membrane footprint of N-BAR, Amph-FL, and Amph CTD ∆SH3 monomers were assumed to be: 16.5 (Adam et al, Scientific Reports 2015) (Adam et al., 2015), 79, and 79 nm^2^, respectively. The average membrane coverage within an experiment was estimated as the mean of all individual vesicle coverage values. Experiments were repeated three times for each protein concentration. Fig. S2D,E and Fig. 3D plots the mean ± first s.d. of the coverage values from the three experiments. To confirm the validity of this analysis approach, we also plotted the area of membrane-bound proteins as a function of vesicle surface area, as shown in Fig. S2M for the 1 μM Amph CTD ∆SH3 dataset. The slope of a linear fit to these data provides an alternative estimate of membrane coverage. The slope of 0.21, or 21% coverage, agrees well with 24% membrane coverage in Fig. 3D that was estimated using the method described above.

Fig. S2D,E plots membrane coverage of N-BAR and Amph-FL at 5, 10, and 25 nM, indicating similar binding behavior of each protein within this concentration regime. At concentrations above 25 nM, Amph-FL significantly deformed and tubulated 200 nm tethered vesicles, leading to inaccuracies in coverage estimates. Therefore, to obtain estimates of membrane coverage by Amph-FL at higher concentrations at which potent fission occurs, we used higher initial curvature vesicles, extruded to 30 nm. Fig. S2F plots membrane coverage by Amph-FL at 25 and 100 nM on 30 nm vesicles, showing that Amph-FL reached approximately 77% membrane coverage at 100 nM, significantly higher than can be reached by protein monomers that do not assemble (Feder, 1980). This coverage is expected to generate significant steric pressure from disordered domain crowding, thus providing a potential explanation for the strong membrane fission observed with Amph-FL at 100 nM.

### Generation of BFP-CLC RPE cell line

A plasmid for expression of BFP-tagged clathrin light chain (BFP-CLC) was generated by replacing the mCherry domain of mCherry-CLC, a gift from Dr. Tom Kirchhausen (Addgene #53972). The mCherry fluorophore was removed and replaced with tagBFP, a gift from Dr. Franck Perez (Addgene #65257). mCherry was excised from the mCherry-CLC plasmid using AgeI and XhoI restriction enzymes. TagBFP was amplified from the li-Str_ManII-SBP-tagBFP plasmid using PCR primers which introduced AgeI and XhoI restriction sites. The resulting tagBFP sequence was digested and ligated onto the CLC backbone to generate BFP-CLC with a linker sequence of HKGRPTR. The CLC-BFP construct was then excised using AgeI and EcoRI restriction sites and ligated into a pLJM1 backbone obtained from Addgene as a gift from Dr. David Sabatini (Addgene #19319). Once subcloned into this viral transfer plasmid, lentiviruses were generated by transfecting the BFP-CLC construct with the envelope plasmid VSVG (a gift from Dr. Jennifer Lippincott-Schwartz, Addgene #11912) and packaging plasmid pCMV-dR8.91 (a gift from Dr. Janet Zoldan). Lentiviral particles were then harvested, filtered, and incubated with human retinal pigmented epithelial (RPE) recipient cells (ARPE-19, purchased from American Type Culture Collection). Cells were incubated with 2 μg/mL puromycin for one week to select for transduced cells which were then used to generate the monoclonal cell line stably expressing BFP-CLC.

### Cell culture and transfection

BFP-CLC RPE cells were grown in 1:1 F12:DMEM supplemented with 10% FBS, 20 mM HEPES, Pen/Strep/L-glutamine (100 units/ml, 100 μg/ml, 300 μg/ml respectively) and incubated at 37°C with 5% CO_2_. Cells were seeded onto acid washed coverslips at a density of 5 × 10^4^ cells per coverslip for 24 hours before transfection with 1 −2 μg of plasmid DNA using 3 μL Fugene transfection reagent per μg of DNA (Promega, Madison, WI, USA). Cells were imaged 16-20 h after transfection. Two independent transfections were performed for each plasmid construct, and data were pooled from both transfections.

Spinning disc confocal z-stacks of BFP-CLC and the mCherry fusion protein were collected with a z-step of 0.25 μm. Z-stacks were analyzed for the number of tubes per cell and tube length. Image analysis was performed using ImageJ software. The plasma membrane frame was chosen by identifying the BFP-CLC frame in which the clathrin-coated structures were best in focus. The plasma membrane expression level of the mCherry fusion protein was then quantified by measuring the mean brightness on a region of the plasma membrane, away from the nucleus and bright structures. Membrane tubes were counted at one frame above the plasma membrane frame. Tube lengths were quantified as the end-to-end distances of the tubes.

Fig. S4A shows an image of a cell stained with CellMask Green plasma membrane stain (Thermo Fisher). Before imaging, the cells were incubated for 5 min at 37 °C in a solution of the CellMask Green stain diluted 1000-fold in sterile PBS. The solution was removed and the cells were washed three times with media before imaging.

A custom-built TIRF microscope was used to collect time-lapse movies of live cells. A 532 nm laser was used to excite mCherry, and a 635 nm laser was used for autofocus. An Olympus IX73 microscope body was equipped with a Photometrics Evolve Delta EMCCD camera and a Zeiss plan-apochromat 100 × 1.46 NA oil immersion TIRF objective. The objective was heated to 37 °C using a Pecon TempController 2000-2 objective heater. The emission filter for the 532 nm laser was a dual bandpass filter centered at 583 nm with 37 nm width and 707 nm with 51 nm width, which minimized signal from the autofocus laser. Movies were collected at the plasma membrane just above the coverslip surface in 2 s intervals for 120 frames. Tube lifetimes and intensities were quantified from TIRF movies. Only movies of cells with similar expression level, acquired under identical imaging settings, were used for analysis. For the tube lifetime analysis in Fig. 4F, only tubes which appeared within the time course of imaging and departed before the end of the time course were included. For the tube intensity analysis, a single frame in the movie with the maximum number of tubes was selected, and the average tube intensity was measured along a straight line drawn on the tube. The mean intensity along an identical line on either side of the tube was also measured, and these values were averaged to estimate the local background intensity of the tube. The protein enrichment on the tube was then quantified as the ratio of the tube intensity to the local background, after subtracting the camera noise background from both values.

### Statistics and sample sizes

TEM experiments: Vesicle diameter distributions in Fig. 1E (Amph-FL) are composed of n > 1,000 vesicles for each condition. Tubule diameter distributions in Fig. S1C (N-BAR) and S5B (F-BAR) are composed of n > 300 and n > 500 tubules, respectively. Vesicle diameter distributions in Fig. S5D (FCHo1-FL) are composed of n > 1,300 vesicles for each condition.

SUPER template experiments: Plots in Fig. 1F, 3H, and 5C display n = 3 independent measurements of SUPER template membrane release at each protein concentration. The indicated p values were calculated using unpaired, one-tailed Student’s t-tests.

Tethered vesicle fission experiments: Vesicle diameter distributions in Fig. 2C-E, 3B,F, 5D,I,J, S3B-D, and S5E represent data pooled from three independent experiments at each protein concentration. Fig. 2C-E (Amph-FL and N-BAR): n > 3,500 vesicles for each condition. Fig. 3B (Amph CTD ∆SH3): n > 4,100. Fig. 3F (N-BAR-epsin CTD): n > 1,000. Fig. 5D (I-BAR-AP180 CTD): n > 4,800. Fig. 5I,J (FCHo1-FL and F-BAR): n > 900. Fig. S3B-D (Amph-FL and N-BAR on highly charged vesicles): n > 800. Fig. S5E (FCHo1-FL on DPPC vesicles): n > 3,900. The markers in Fig. 2F, 3C,G, 5K, S3E, and S5G show the mean ± first s.d. of the three independent experiments.

Protein coverage experiments on tethered vesicles: Markers in Fig. S2D,E (Amph-FL and N-BAR) and Fig. 3D (Amph CTD ∆SH3) show the mean ± first s.d. from three independent experiments, where n > 4,100 total vesicles for each concentration in Fig. S2D,E and n > 2,900 total vesicles for each concentration in Fig. 3D. Fig. S2F (Amph-FL on 30 nm extruded vesicles) show data from one experiment where n > 1,700 vesicles for each concentration. Bars show mean ± 95% confidence interval.

Cell experiments: Fig. 4C displays data from n > 90 cells per condition from two independent transfections. Fig. 4D displays a subset of the data in Fig. 4C that is within the specified protein expression range, with n > 20 cells per condition. Bars represent mean ± s.e.m. Fig. 4E displays the lengths of individual tubes from cells within the specified protein expression range, where n > 80 tubes per condition. Fig. 4F displays the lifetimes of individual tubes measured from TIRF movies, where n > 40 tubes per condition. Black lines in Fig. 4E,F indicate means. The indicated p-values were calculated using unpaired, two-tailed Student’s t-tests.

### Calculation of IDP compression above BAR scaffold

The volume per disordered domain (IDP) attached to the BAR scaffold was estimated as the volume of a cylindrical shell surrounding a membrane tube, with thickness equal to twice the radius of gyration of the IDP domains, divided by the number of BAR domains in the scaffold. The cylindrical shell volume surrounding the membrane tube is therefore:

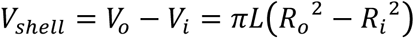

Where *V_o_* and *V_i_* are the outer and inner radii of the shell, respectively, *L* is the tube length, *R_i_* is the radius of the membrane tube, and *R_o_ = R_i_ + 2r_IDP_*, with *r_IDP_* equal to the radius of gyration of amphiphysin’s disordered domain. The number of proteins in the scaffold is:

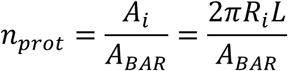

Where *A_i_* is the surface area of the membrane tube and *A_BAR_* is the area occupied per BAR monomer. The volume per compressed, scaffold-anchored disordered domain is:

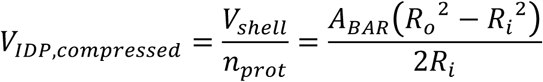
and the un-compressed volume of the disordered domain is:

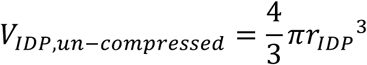

Parameter values were taken as: *A_BAR_ =* 16.5 *nm*^2^*, R_i_ =* 14 *nm* (both from Adam et al, Scientific Reports 2015) (Adam et al., 2015), and *r_IDP_ =* 5*nm*. Using these values, *V_IDP,compressed_ =* 224 *nm*^3^ and *V_IDP,un–compressed_ =* 524 *nm*^3^, corresponding to an approximately 60% compression of the disordered domain volume to accommodate the scaffold geometry. See also Fig. 3I.

## Acknowledgements

J.C.S. and E.M.L. acknowledge funding from the National Institutes of Health (R01GM112065), including an administrative supplement for diversity in support of W.F.Z. W.T.S. acknowledges the support of a Ruth L. Kirschstein NRSA Predoctoral Fellowship from the National Institutes of Health (F31GM121013), as well as a fellowship from the Graduate School at UT Austin. G.K. acknowledges the support of a National Science Foundation Graduate Research Fellowship. We thank Dr. Carl Hayden for assistance with fluorescence microscopy and FCS experiments. We thank Dr. Anjon Audhya (University of Wisconsin - Madison) for advice on the purification of full-length FCHo1. We thank Liping Wang (Lafer lab, UT Health Science Center at San Antonio) for assistance with protein expression and purification. We thank Dr. Darryl Sasaki (Sandia National Laboratories, Livermore, CA) for providing DP-EG10-biotin. We thank Dr. Gaudenz Danuser and Dr. Sandra Schmid (UT Southwestern, Dallas, TX) for freely providing cmeAnalysis particle detection software. We thank Dr. Dwight Romanovicz and the ICMB Microscopy Facility at UT Austin for assistance with electron microscopy. The authors declare no competing financial interests.

## Author Contributions

All authors designed and performed experiments, consulted together on the data, and wrote the manuscript.

## Supplemental Material

**Figure S1.**
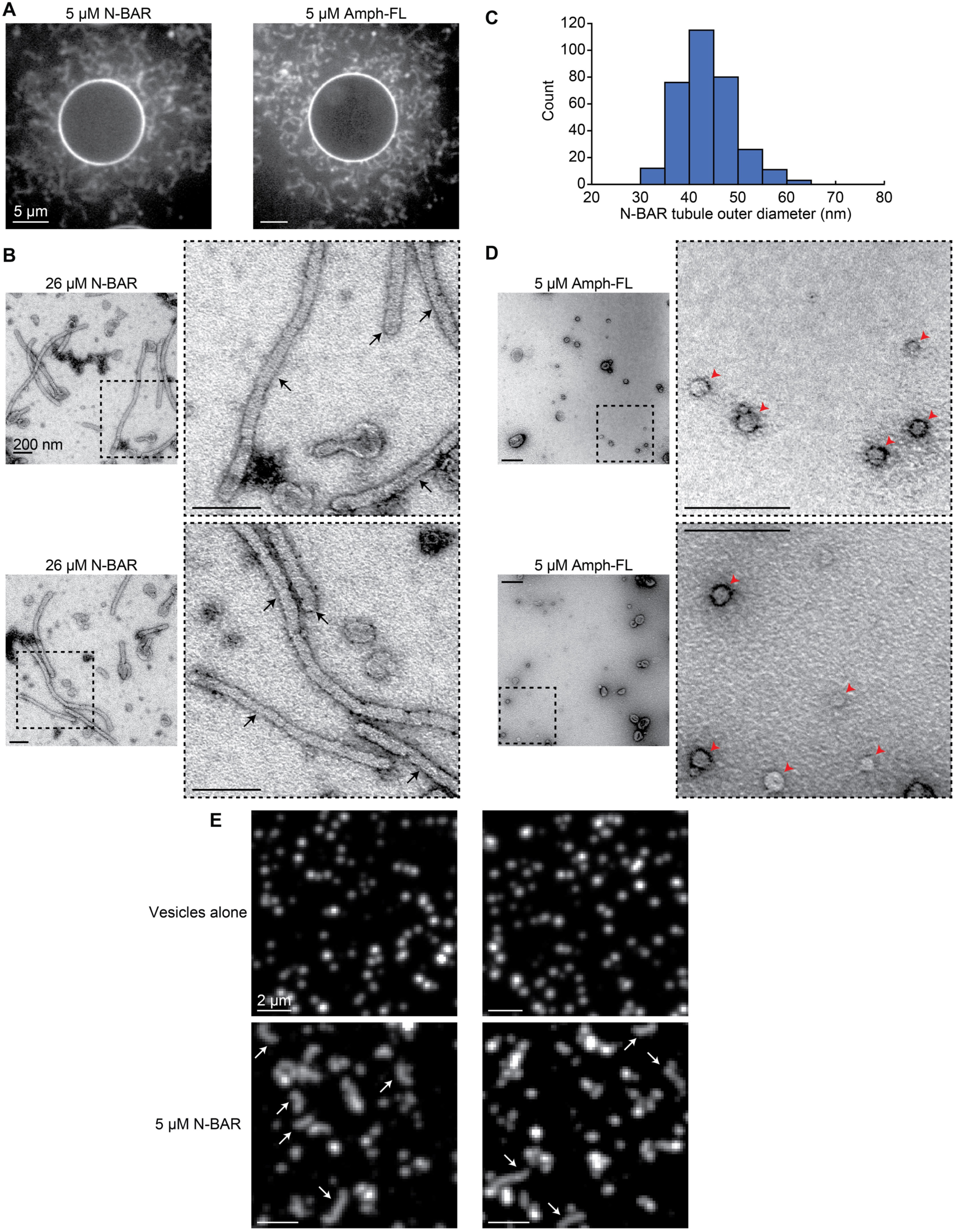
While N-BAR generates membrane tubules, Amph-FL forms highly curved fission vesicles. GUV membrane composition: 79.5 mol% DOPC, 5 mol% PtdIns(4,5)P_2_, 15 mol% DOPS, 0.5 mol% Oregon Green 488-DHPE. Membrane composition in TEM experiments: 80 mol% DOPC, 5 mol% PtdIns(4,5)P_2_, 15 mol% DOPS, extruded to 200 nm. Membrane composition in tethered vesicle experiments: 76 mol% DOPC, 5 mol% PtdIns(4,5)P_2_, 15 mol% DOPS, 2 mol% Oregon Green 488-DHPE, 2 mol% DP-EG10-biotin, extruded to 200 nm. (**A**) Representative spinning disc confocal micrographs of GUVs after exposure to 5 μM of either N-BAR (left) or Amph-FL (right). Fluorescence signal comes from Atto 594-labeled protein. Scale bars: 5 μm. (**B**) Two representative electron micrographs of tubules generated by 26 μM N-BAR. Dashed boxes indicate zoomed regions to the right of each image. Black arrows indicate tubules. Scale bars, including zoomed regions: 200 nm. (**C**) Histogram of tubule outer diameters generated by 26 μM N-BAR. Mean = 44 ± 6 nm first s.d., n = 323 tubules. (**D**) Two representative electron micrographs of fission vesicles generated by 5 μM Amph-FL. Dashed boxes indicate zoomed regions to the right of each image. Red arrowheads indicate fission vesicles. Scale bars, including zoomed regions: 200 nm. (**E**) Two representative spinning disc confocal micrographs of tethered vesicles before exposure to protein (top) and after exposure to 5 μM N-BAR (bottom). At 5 μM, a higher concentration than used in experiments in Fig. 2, N-BAR generates membrane tubules with length greater than the diffraction limit of light. Tubules indicated by white arrows. Scale bars: 2 μm.

**Figure S2.**
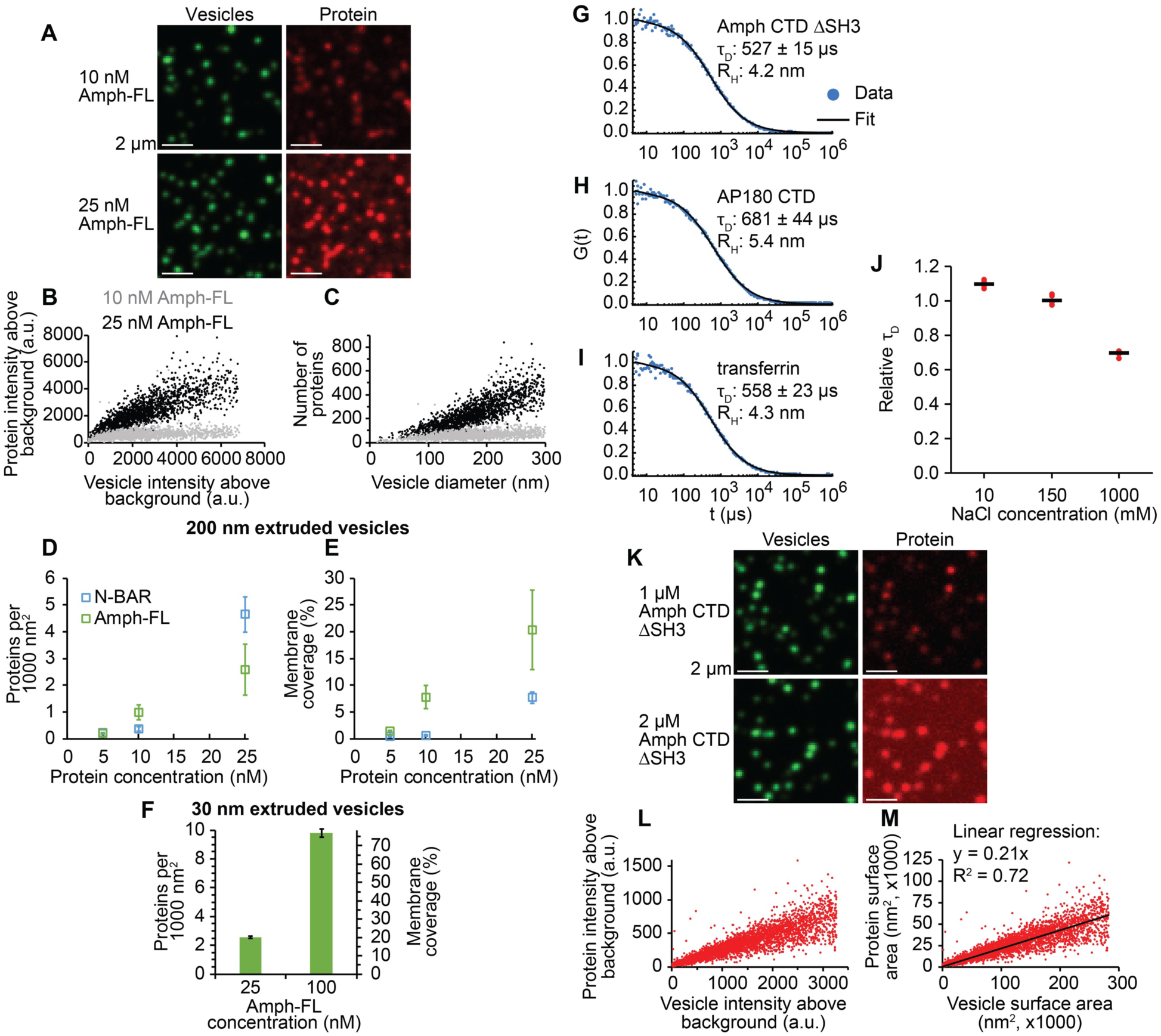
Membrane coverage measurements of N-BAR, Amph-FL, and Amph CTD ∆SH3, and fluorescence correlation spectroscopy (FCS) of Amph CTD ∆SH3. Tethered vesicle composition for N-BAR and Amph-FL: 76 mol% DOPC, 15 mol% DOPS, 5 mol% PtdIns(4,5)P2, 2 mol% DP-EG10-biotin, and 2 mol% Oregon Green 488-DHPE. In experiments with Amph CTD ∆SH3, DOPS and PtdIns(4,5)P_2_ were replaced by 20 mol% DOGS-NTA-Ni. (A-E and K-M): Extruded to 200 nm. (F): Extruded to 30 nm. (**A**) Images of tethered vesicles (green, left column) and membrane-bound Amph-FL-Atto 594 (red, right column). Top row is 10 nM, bottom row is 25 nM. Images in each column have equal contrast to show that increasing protein concentration leads to increased membrane-bound protein intensity. Scale bars: 2 μm. (**B**) Raw protein intensity as a function of raw vesicle intensity for 10 and 25 nM Amph-FL. (**C**) The same data in (B) after processing (see methods), plotted as the number of membrane-bound proteins as a function of vesicle diameter. (**D**) Number of membrane-bound proteins per 1000 nm^2^ of membrane surface area as a function of the concentration of N-BAR or Amph-FL. The two proteins follow a similar trend, indicating similar binding. (**E**) Data in (D) plotted as the coverage of the membrane surface by proteins as a function of protein concentration. Because Amph-FL occupies a greater area on the membrane surface compared to N-BAR (79 versus 16.5 nm^2^ per monomer, respectively), Amph-FL reaches a higher coverage than N-BAR at equal number density of membrane-bound proteins. At 25 nM, Amph-FL reaches approximately 20% membrane coverage, approaching a crowded regime (Snead et al., 2017; Stachowiak et al., 2012). Markers in (D) and (E) represent mean ± first s.d., n = 3 independent experiments. (**F**) Membrane coverage estimates for 25 and 100 nM Amph-FL on 30 nm vesicles. At 100 nM Amph-FL, when potent membrane fission occurs, Amph-FL reaches approximately 77% membrane coverage, significantly higher than can be reached by non-assembling proteins. At this coverage, steric pressure from disordered domain crowding is very high, providing a potential explanation for strong membrane fission by Amph-FL. Markers indicate mean of all coverage values, error bars represent 95% confidence interval. 25 nM: n = 2,171 vesicles, 100 nM: n = 1,783 vesicles. (**G-I**) Representative, normalized FCS traces of (**G**) Amph CTD ∆SH3, (**H**) AP180 CTD, and (**I**) transferrin. Blue dots indicate data, black lines indicate fit (see methods). Average values of diffusion time, τ_D_, ± first s.d. are shown next to each trace, with n = 10, 5, and 3 FCS traces for Amph CTD ∆SH3, AP180 CTD, and transferrin, respectively. The hydrodynamic radius, *R_H_*, of each protein is also shown next to each trace. *R_H_* of Amph CTD ∆SH3 was computed by scaling from the known *R_H_* of AP180 CTD (Busch et al., 2015) (see methods). Using this same approach yields an *R_H_* of transferrin that is similar to its expected value (Hall et al., 2002). (**J**) Relative diffusion times of Amph CTD ∆SH3 in buffer with 10, 150, and 1,000 mM sodium chloride (NaCl), expressed as the proportion of the average diffusion time at 150 mM NaCl. The data indicate a transition from an extended to a more compact state with increasing ionic strength, as expected for charged disordered proteins (Srinivasan et al., 2014). Diffusion times were corrected for changes in solution viscosity with varying NaCl concentration (Zhang and Han, 1996). Red dots indicate data, black lines indicate mean. n = 5 FCS traces at each NaCl concentration. (**K**) Images of tethered vesicles (green, left column) and membrane-bound Amph CTD ∆SH3-Atto 594 (red, right column). Top row is 1 μM, bottom row is 2 μM. Images in each column have equal contrast to show greater protein intensity with increasing concentration. Scale bars: 2 μm. (**L**) Raw protein intensity as a function of raw vesicle intensity for the 1 μM Amph CTD ∆SH3 dataset. (**M**) The same 1 μM Amph CTD ∆SH3 dataset after processing, plotted as the area occupied by membrane-bound proteins as a function of vesicle surface area. The slope of a linear fit to the data represents the average membrane coverage by proteins. The value of 21% coverage from this approach agrees with 24% membrane coverage shown in Fig. 3D, estimated by averaging all of the membrane coverage values of individual vesicles.

**Figure S3.**
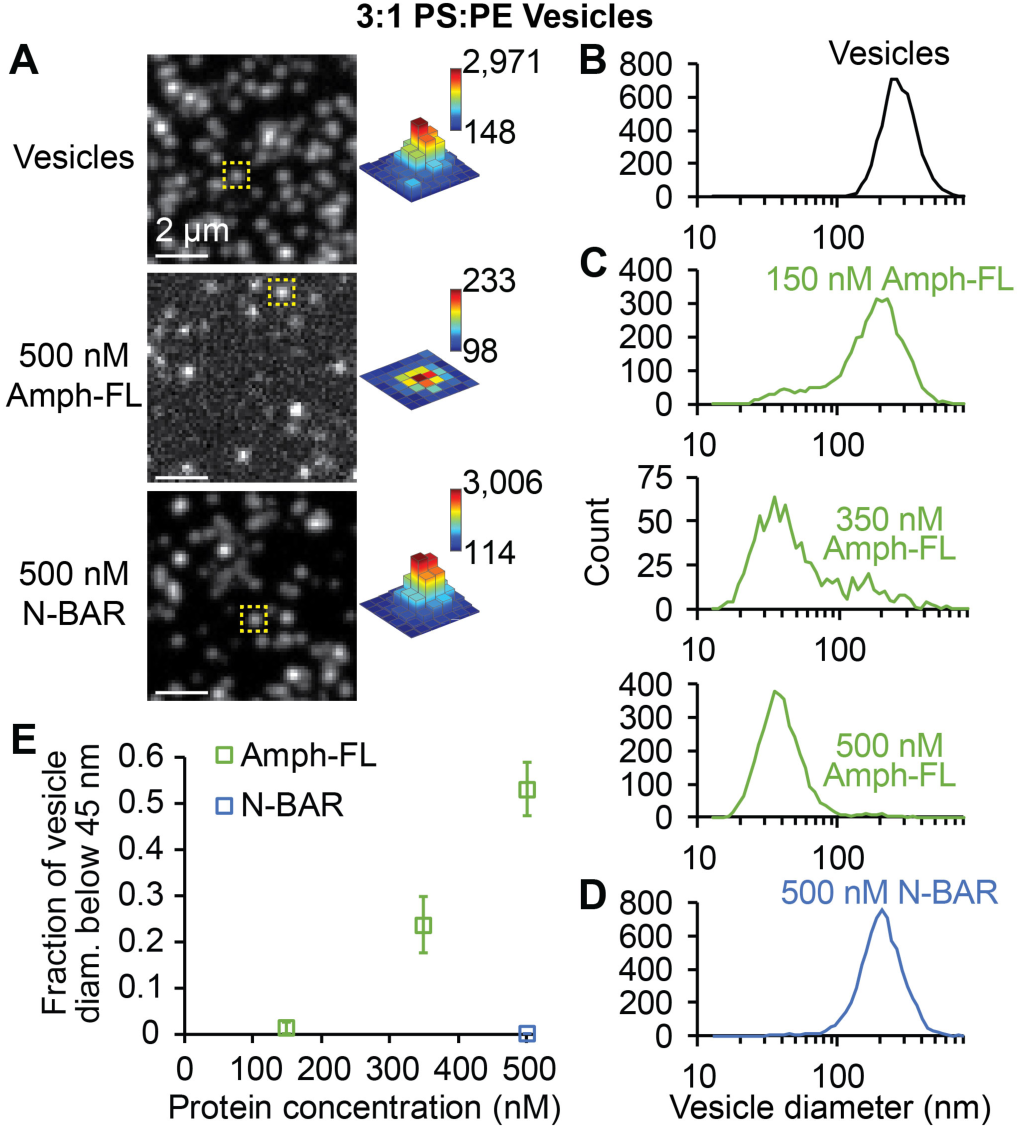
Amph-FL drives fission of membranes that contain high concentrations of negatively charged lipids. Vesicle composition: 68 mol% DOPS, 23 mol% DOPE, 5 mol% cholesterol, 2 mol% DP-EG10-biotin, and 2 mol% Oregon Green 488-DHPE, extruded to 200 nm. Composition based on Mim et al, Cell 2012 (Mim et al., 2012). (**A**) Representative spinning disc confocal micrographs of vesicles (top) before protein exposure, (middle) after exposure to 500 nM Amph-FL, and (bottom) after exposure to 500 nM N-BAR. Contrast settings in top and bottom images are the same while contrast in middle image is adjusted to clearly show vesicle puncta. Dashed yellow boxes indicate puncta intensity profiles on right, where bar heights are all scaled between 90 and 3,050 brightness units while each color map corresponds to specified intensity range. Scale bars: 2 μm. (**B-D**) Distributions of vesicle diameter measured by tethered vesicle assay (**B**) before exposure to protein, (**C**) after exposure to Amph-FL at 150, 350, and 500 nM, and (**D**) after exposure to N-BAR at 500 nM. (**E**) Summary of tethered vesicle fission data, expressed as the ratio of the distribution area below 45 nm diameter to the total distribution area. Compare to Fig. 2F. Markers represent mean ± first s.d., n = 3 independent experiments.

**Figure S4.**
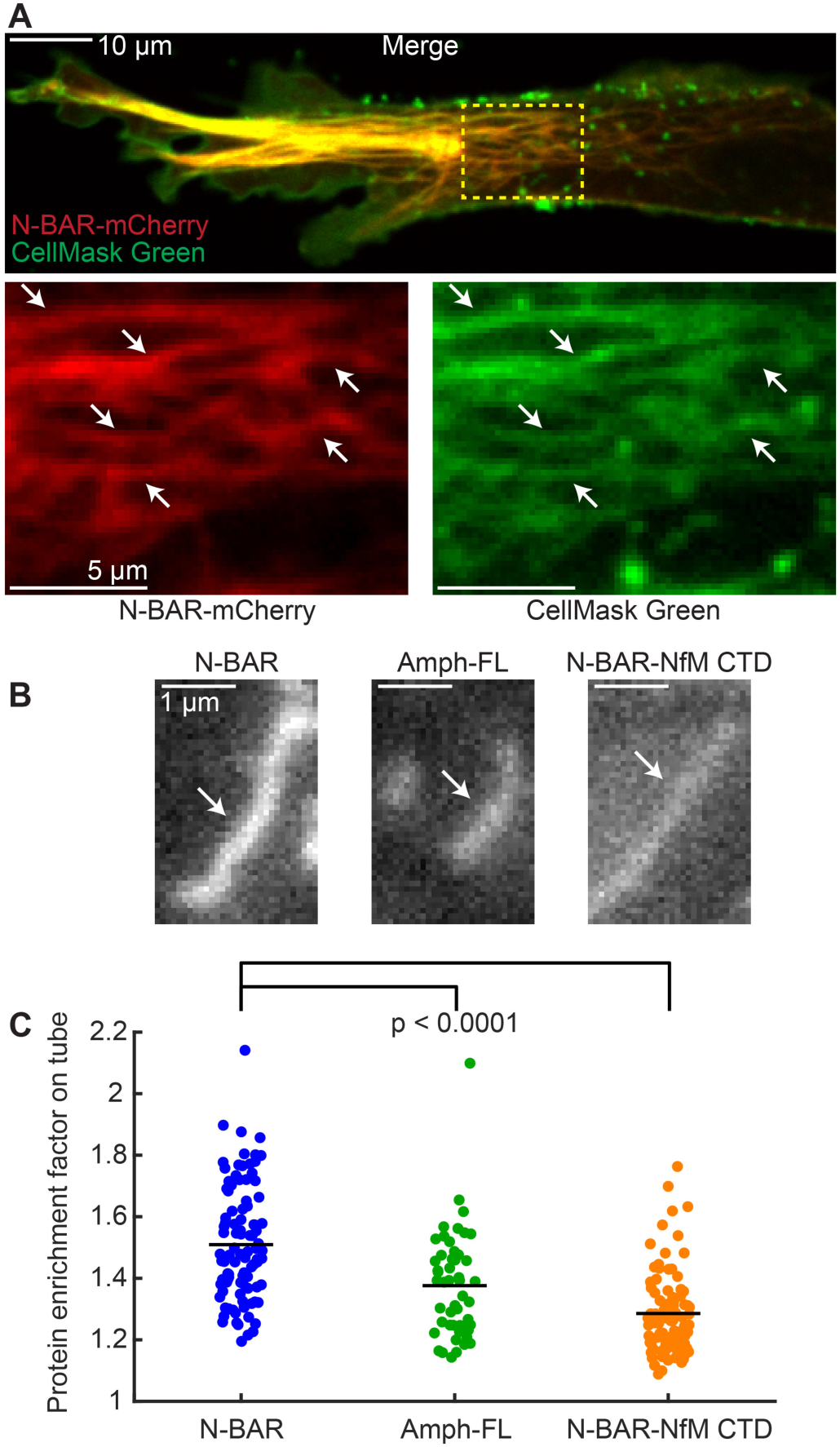
Imaging tubules generated in live cells by N-BAR, Amph-FL, and N-BAR-NfM CTD. (**A**) Spinning disc confocal image of a live RPE cell overexpressing the N-BAR domain of amphiphysin tagged with mCherry and stained with CellMask Green plasma membrane stain. Yellow dashed box indicates the zoomed region below, where the individual channels are shown. White arrows indicate lipid tubes. The co-localization of the plasma membrane stain with N-BAR-coated tubes indicates that the tubes are derived from the plasma membrane. Scale bar in top image: 10 μm. Scale bar in zoomed region: 5 μm. (**B**) Representative tubes in RPE cells overexpressing the indicated proteins tagged with mCherry. Images were acquired in TIRF at 37 °C. Tubules are from cells with similar protein expression level, and all images are displayed with equal contrast settings. White arrows indicate tubes. N-BAR shows greater enrichment on the tube relative to the local background compared to Amph-FL and N-BAR-NfM CTD. Scale bars: 1 μm. (**C**) Protein intensity on membrane tubes in live cells, quantified as the ratio of the tube intensity to the local background intensity. Points indicate individual tubes, and black lines indicate means. Data were quantified from TIRF movies that were taken under identical imaging settings. n = 100, 50, and 91 tubes for N-BAR, Amph-FL, and N-BAR-NfM CTD, respectively. P-values: unpaired, two-tailed Student’s t-tests. These data indicate the disordered domains of Amph-FL and N-BAR-NfM CTD did not promote fission of lipid tubules in live cells by enhancing protein binding to the membrane surface.

**Figure S5.**
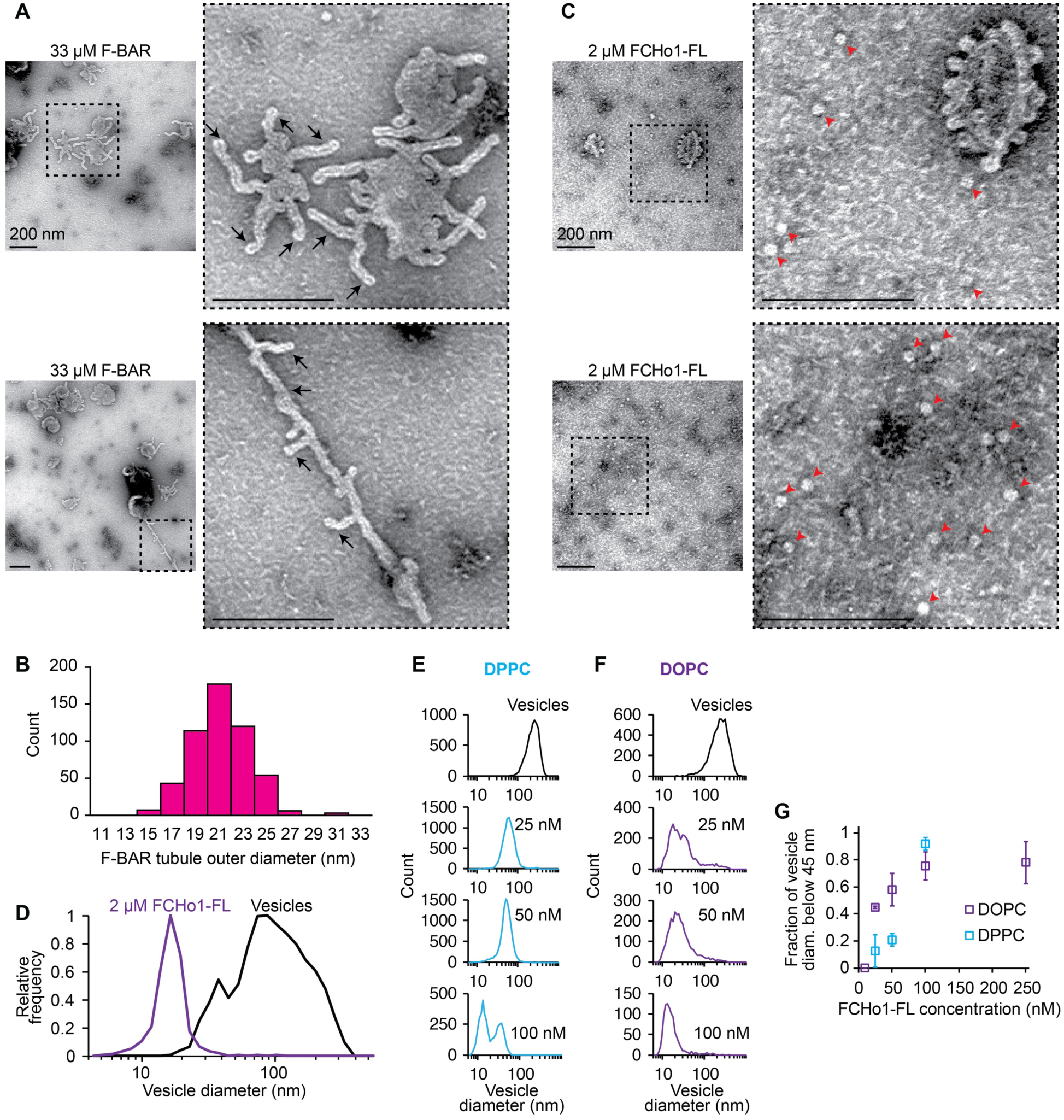
While F-BAR generates membrane tubules, FCHo1-FL forms highly curved fission vesicles. Membrane composition in TEM experiments: 80 mol% DOPC, 5 mol% PtdIns(4,5)P_2_, 15 mol% DOPS. Vesicle composition in (E): 76 mol% DPPC, 15 mol% DPPS, 5 mol% PtdIns(4,5)P_2_, 2 mol% DP-EG10-biotin, and 2 mol% Oregon Green 488-DHPE. For vesicles in (F), DPPC and DPPS were replaced with DOPC and DOPS, respectively. All vesicles extruded to 200 nm. (**A**) Two representative electron micrographs of tubules generated by 33 μM F-BAR. Dashed boxes indicate zoomed regions to the right of each image. Black arrows indicate tubules. Scale bars, including zoomed regions: 200 nm. (**B**) Histogram of the outer diameters of tubules generated by 33 μM F-BAR. Mean = 21 ± 2 nm first s.d., n = 524 tubules. (**C**) Two representative electron micrographs of fission vesicles generated by 2 μM FCHo1-FL. Dashed boxes indicate zoomed regions to the right of each image. The top zoomed region shows a larger vesicle with highly curved buds extending from the surface, which may be an intermediate state prior to full fission. Red arrowheads indicate fission vesicles. Scale bars, including zoomed regions: 200 nm. (**D**) Histograms of vesicle diameters before and after exposure to 2 μM FCHo1-FL. FCHo1-FL generates a high-curvature vesicle population with average diameter 17 ± 7 nm first s.d. Vesicles alone: n = 1,302 vesicles. 2 μM FCHo1-FL: n = 1,676 vesicles. (**E**) Fission product diameter distributions from tethered vesicle fission assay with DPPC as the primary lipid after exposure to 25, 50, and 100 nM FCHo1-FL. (**F**) Fission product diameter distributions with DOPC as the primary lipid after 25, 50, and 100 nM FCHo1-FL, repeated from Fig. 5I. DPPC fission product distributions in (E) show lower-curvature populations compared to the high-curvature populations with DOPC vesicles in (F), indicating that increasing the bilayer rigidity restricts membrane fission by FCHo1-FL. (**G**) Plot of the proportion of fission product diameters below 45 nm as a function of FCHo1-FL concentration. DPPC vesicles require higher FCHo1-FL concentration to observe high-curvature fission products observed with DOPC vesicles. Markers represent mean ± first s.d., n = 3 independent experiments. DOPC data repeated from Fig. 5K.

## Supplemental Movie Legends

**Movie S1.** The amphiphysin N-BAR domain generates mobile lipid tubules from GUVs. GUV composition: 79.5 mol% DOPC, 5 mol% PtdIns(4,5)P_2_, 15 mol% DOPS, 0.5 mol% Oregon Green 488-DHPE. GUVs were mixed with 5 μM N-BAR and imaged by confocal microscopy. Fluorescence signal comes from Atto594-labeled protein. The frames are approximately 400 ms apart. The video plays at 5 frames per second.

**Movie S2.** Full-length amphiphysin generates mobile lipid tubules from GUVs. GUV composition: 79.5 mol% DOPC, 5 mol% PtdIns(4,5)P_2_, 15 mol% DOPS, 0.5 mol% Oregon Green 488-DHPE. GUVs were mixed with 5 μM Amph-FL and imaged by confocal microscopy. Fluorescence signal comes from Atto594-labeled protein. The frames are approximately 400 ms apart. The video plays at 5 frames per second.

**Movie S3.** The amphiphysin N-BAR domain drives collapsing of vesicles into diffraction-limited tubules and fragments. GUV composition: 79.5 mol% DOPC, 5 mol% PtdIns(4,5)P_2_, 15 mol% DOPS, 0.5 mol% Oregon Green 488-DHPE. GUVs were mixed with 5 μM N-BAR and imaged by confocal microscopy. Fluorescence signal comes from Atto594-labeled protein. The frames are approximately 400 ms apart. The video plays at 5 frames per second.

**Movie S4.** Full-length amphiphysin drives collapsing of vesicles into diffraction-limited tubules and fragments. GUV composition: 79.5 mol% DOPC, 5 mol% PtdIns(4,5)P_2_, 15 mol% DOPS, 0.5 mol% Oregon Green 488-DHPE. GUVs were mixed with 5 μM Amph-FL and imaged by confocal microscopy. Fluorescence signal comes from Atto594-labeled protein. The frames are approximately 400 ms apart. The video plays at 5 frames per second.

**Movie S5.** Live cell imaging reveals that N-BAR generates tubules with longer lifetime compared to Amph-FL and N-BAR-NfM CTD. Movie shows tubules in live RPE cells expressing N-BAR (left), Amph-FL (middle), or N-BAR-NfM CTD (right), imaged by TIRF microscopy at 37 °C. The cells are all at similar protein expression level, and were imaged using identical settings. The N-BAR-expressing cell shows a greater number of tubules that also persist longer compared to Amph-FL and N-BAR-NfM CTD. See Fig. 4F for quantification. The frames are 2 s apart, 120 frames total. The video plays as 10 frames per second.

**Movie S6.** The IRSp53 I-BAR domain drives inward tubulation. GUV composition: 79.5 mol% DOPC, 5 mol% PtdIns(4,5)P2, 15 mol% DOPS, 0.5 mol% Oregon Green 488-DHPE. GUVs were mixed with 5 μM I-BAR and imaged by confocal microscopy. Fluorescence signal comes from Atto594-labeled protein. The frames are approximately 500 ms apart. The video plays at 5 frames per second.

**Movie S7.** The I-BAR-AP180 CTD chimera drives frustrated membrane fluctuations. GUV composition: 79.5 mol% DOPC, 5 mol% PtdIns(4,5)P_2_, 15 mol% DOPS, 0.5 mol% Oregon Green 488-DHPE. GUVs were mixed with 10 μM I-BAR-AP180 CTD and imaged by confocal microscopy. Fluorescence signal comes from Atto594-labeled protein. The frames are approximately 500 ms apart. The video plays at 5 frames per second.

